# Predictive coding of natural images by V1 activity revealed by self-supervised deep neural networks

**DOI:** 10.1101/2020.08.10.242958

**Authors:** Cem Uran, Alina Peter, Andreea Lazar, William Barnes, Johanna Klon-Lipok, Katharine A Shapcott, Rasmus Roese, Pascal Fries, Wolf Singer, Martin Vinck

**Author notes:** Equally contributing (Cem Uran, Alina Peter).

## Abstract

Predictive coding is an important candidate theory of self-supervised learning in the brain. Its central idea is that neural activity results from an integration and comparison of bottom-up inputs with contextual predictions, a process in which firing rates and synchronization may play distinct roles. Here, we quantified stimulus predictability for natural images based on self-supervised, generative neural networks. When the precise pixel structure of a stimulus falling into the V1 receptive field (RF) was predicted by the spatial context, V1 exhibited characteristic *γ*-synchronization (30-80Hz), despite no detectable modulation of firing rates. In contrast to *γ, β*-synchronization emerged exclusively for unpredictable stimuli. Natural images with high structural predictability were characterized by high compressibility and low dimensionality. Yet, perceptual similarity was mainly determined by higher-level features of natural stimuli, not by the precise pixel structure. When higher-level features of the stimulus in the receptive field were predicted by the context, neurons showed a strong reduction in firing rates and an increase in surround suppression that was dissociated from synchronization patterns. These findings reveal distinct roles of synchronization and firing rates in the predictive coding of natural images.

## Introduction

The ability to make accurate predictions is central to adaptive behavior in a complex world, and relies on the existence of strong regularities in both time and space. For example, when we look at a normal face, we recognize that the elements (mouth, nose, etc.) are in the expected spatial configuration. But, if we were to see a scrambled face with the mouth above the eyes, we would be surprised and explore whether we had been tricked or had discovered a new facial configuration. The brain can perform these kind of predictive operations by comparing sensory inputs with the spatiotemporal context in which they are embedded. Predictive signalling may be important for a host of sensory processes: It can shape perception, as illustrated by perceptual Gestalt principles and illusions; it can facilitate efficient encoding of information (1; 2; 3); it can also provide a mechanism by which the brain tests hypotheses about incoming sensory data and performs inference (1; 4; 3); finally, predictive signaling can play a role in self-supervised learning, i.e. learning based on the data itself by predicting a subset of inputs based on another subset of inputs, without requiring human labels (5; 6; 7). Despite predictive signaling’s many postulated roles, a detailed understanding has yet to be reached of how neuronal signals are precisely modulated by the relationship between bottom-up inputs and contextual predictions during natural vision.

In canonical feedforward models of visual processing, response properties of neurons are described based on their classical receptive field (RF) in a circumscribed region of visual space (8). However, processing abstracted stimuli through punctual activation of the RF is not the case when the active brain encounters natural stimuli. That is, natural RF stimuli are embedded in a spatiotemporal context with certain statistical regularities (9). Stimuli presented outside the classical RF can strongly influence responses in visual areas like V1 (10; 11; 12; 13; 14; 15). Non classical RF influences likely depend both on horizontal and top-down feedback (16; 17), and can take the form of sparsification, suppression (13; 14), or facilitation (15). Different theories have been proposed about the way in which contextual predictions interact with bottom-up inputs. According to the predictive coding model of (1), V1 output firing rates represent prediction errors that signal the difference between bottom-up inputs into V1 and predictions from the spatio-temporal context. Different flavors of V1 models in which predictive processing plays a role have been also proposed (18; 19; 20; 1; 3). Importantly, the computational and mechanistic underpinnings of the integration between bottom-inputs and contextual predictions remain unclear. Because contextual predictions likely reflect our innate and learned priors about the statistics of the natural environment, it is critical to examine predictive processing using natural scenes rather than artificial stimuli because artificial stimuli may be extreme outliers in our visual environment (e.g., gratings, homogeneous surfaces). However, studying predictive processing for natural images is a fundamentally difficult problem, because the constructs of “prediction” and “prediction error” for natural images need to be mathematically operationalized and quantified. Ideally this would rely on generative neural networks that learn both linear and non-linear natural scene statistics (i.e. priors) across a very large number of images in a self-supervised manner; however such an approach has so far not been used to predict the contextual modulation of V1 activity. Notably, it is also unclear what the precise nature of V1 prediction errors would be. V1 prediction errors can in principle encode sensory predictions or prediction errors of varying levels of complexity, ranging from low-level content (e.g. pixel values) to high-level features (21; 20; 22; 3; 1).

Another feature of V1 activity that may play an important role in predictive processing is *γ*-synchronized firing (23), which reflects balanced interactions between inhibition and excitation (24; 25; 26). Synchronized firing can influence both signal integration and synaptic plasticity induction (27; 28; 29). A number of groups have hypothesized that *γ*-(30-80Hz) and *β*-synchronized (13-30Hz) firing reflects prediction error and prediction signaling, respectively (30; 31; 32; 33). Indirect evidence for this hypothesis comes from the finding that feedforward and feedback V1-V2/V4 communication are particularly strong in the *γ-* and *β*-frequency band, respectively (34; 35), (but see (36)). In contrast, it has been hypothesized that V1 *γ*-synchrony reflects the extent to which RF inputs can be predicted from the surround (37). It is however unclear under which conditions *γ*-synchrony and V1 firing rates signal similar features or convey independent information, as both positive and negative correlations have been reported (38; 37; 39; 40; 41).

Here, we developed a self-supervised DNN (Deep Neural Network) to generate predictions of the likely structure of stimuli falling in a neuron’s RF. These predictions were derived from the spatial context, i.e. the image residing outside of the RF. That is, the network was trained to predict a subset of sensory data (the RF area) based on other subsets of sensory data (the non-RF area). Examining the response properties of nodes at various levels of a CNN (Convolutional Neural Network) for object recognition, we then derived measures of spatial stimulus predictability that distinguished between low- and high-level features. After presenting natural images (naturally varying in spatial stimulus predictability within and between the images) to awake macaque monkeys while recording V1 activity, we analyzed, using the metrics we developed with CNNs, how spatial predictability affects firing rates, synchronization patterns, and surround modulation. Preliminary results of this work have been presented previously at the FENS 2018 and Bernstein 2018 meeting.

## Results

We recorded multi-unit (MU) spiking activity and local field potentials (LFP) from 32-64 channel (semi-)chronic microelectrode arrays in three awake macaque monkeys performing a passive fixation task and viewing large (>11deg) natural images (Figure 1A-C, Figure 2). RF eccentricities were 5.2dva on average (range: 2.5-10.6dva) with an average diameter of 1.44dva (which was likely overestimated because of small eye movements) (Figure 1A). LFP and MUA power spectra showed a typical 1/ *f* trend (Figure 1D, Pesaran et al. (42)) and spectral peaks in the *γ*-frequency band (30-80Hz). These peaks reflect the rhythmic modulation and synchronization of synaptic and spiking activity (Figure 1D-G) (43; 44; 42). We estimated their magnitude by removing the 1/ *f* trend by fitting polynomials as in (45) (see Methods, Figure 1E).

**Figure 1:**
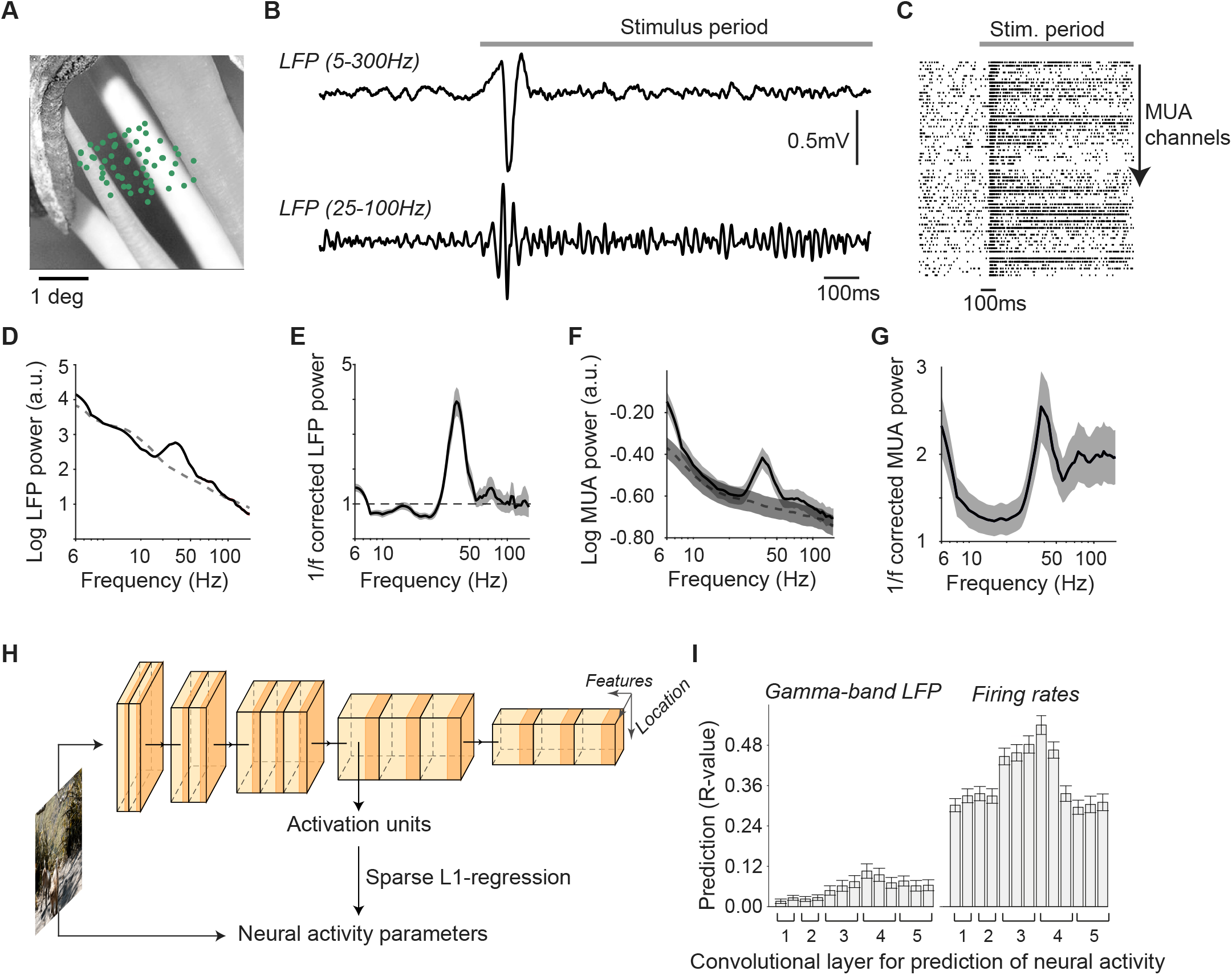
Recording paradigm and prediction of neural activity based on activation of neurons in VGG-16 network for object recognition. **(A)** Natural images were presented for 1.2s (in a subset of sessions, for 0.6s) and subtended 11.5 to 15 degrees of visual angle (dva). Green dots indicate locations of RF centers of the recording array in monkey H. Image is cut out around the RF locations. **B** Local Field Potentials (LFPs) were recorded and showed gamma-band activity after stimulus onset. Shown is a median example trace for the image shown in (A). **C** 64 MUA channels were recorded in each session. Shown is a raster plot (spikes thresholded at 3 s.d). **D** For the example image, LFP spectra showed a characteristic 1/ *f* trend during the baseline period (dashed line), and an increase in gamma-band activity during the stimulus period (solid line). **(E)** After 1/ *f* correction, a clear gamma-peak in the LFP was visible. **(F-G)** MUA spiking activity showed a gamma-peak during visual stimulation both in the raw spectra and in the 1/ *f* - corrected spectrum (solid line). This peak was not present during baseline (dashed line). **H** For each recording site, we determined different neural activity parameters. The image patch centered on the RF of the recording site was then passed into the VGG-16, and we computed the activation of every VGG-16 artificial neuron (AN) whose RF overlapped with the recording site. Sparse L1-regression with cross-validation was used to predict neural activity from VGG-16 ANs with RFs at the center of the image. **(I)** Regression prediction accuracy of different neural activity parameters depending on VGG-16 layer. Regression prediction accuracy for late (200-600 ms) firing rates was significantly higher for middle (5–9) than early (1–4) and deep (10–13) convolutional layers (*P* < 0.001, paired T-test). For *γ*, regression prediction accuracy was significantly higher for middle (*P* < 0.001) and deep (*P* < 0.05) than early layers. For early rates see Figure Extended Data 1.

**Figure 2:**
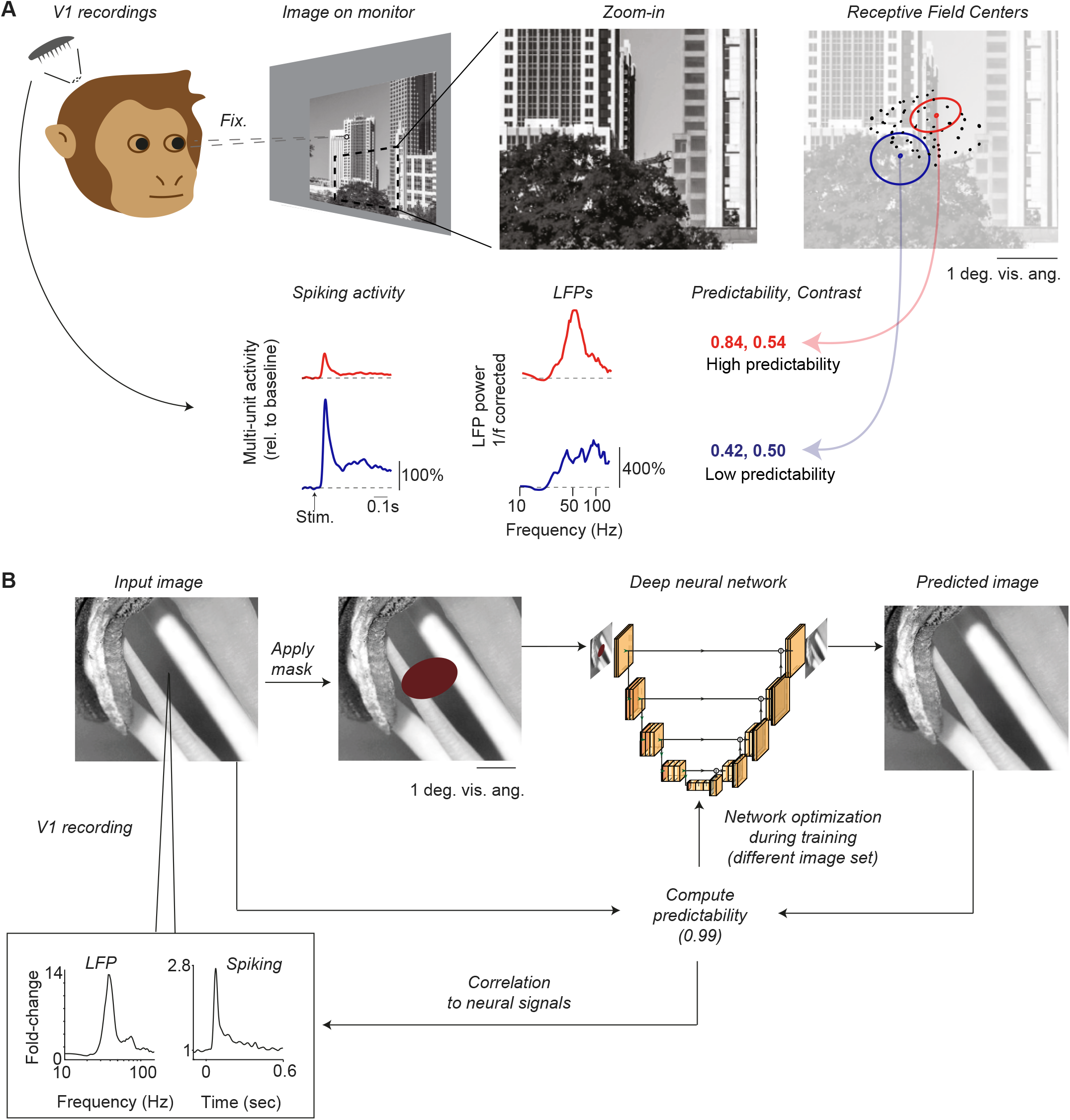
Illustration of experiments and image prediction network. **(A)** We recorded LFPs and spiking activity from area V1 of awake monkeys. Monkeys performed a passive fixation task. Shown are two example sites with their receptive fields (RFs) overlaid on a zoomed-in part of the image. Bottom: left to right: Firing intensity, LFP power spectra, and values of luminance contrast and spatial predictability based on RF locations. The red recording site shows clear narrow-band LFP gamma oscillations around 50 Hz, whereas the blue site shows a broad-band increase in power due to increased spiking activity (41). **(B)** Illustration of a deep neural network trained to predict visual inputs into the RFs. A mask of approximately the same size as the recording site’s RF is applied to an image. The image with the mask is then entered as an input to a deep neural network with a U-net architecture. This network generates/predicts the full image, i.e. the image content behind the mask is filled in. Spatial predictability is computed by comparing the ground-truth input image and the predicted image. The stimulus predictability signal is then used for network optimization during the training stage. After network training, a novel set of images is presented to both the network and the monkeys. We then analyze how spatial predictability determines neural activity. Shown are firing intensity and LFP power for the recording site with a RF corresponding to the mask.

### Predicting neural activity from a feedforward neural network for object recognition

We first investigated to what extent V1 firing rates and synchronization can be explained by a standard feedforward neural network for object recognition, the VGG-16, which is considered to be a state-of-the-art model of the primate ventral stream (Figure 1). This illustrates the performance of standard feed-forward networks in predicting neural activity as a baseline before developing an alternative approach (Figure 2), and introduces the VGG-16 network, which will play an important role later (Figure 6). Convolutional neural networks (CNNs) like the VGG-16 contain multiple layers, with increasingly complex representations in the deepest layers that extract information about object identity. Each layer contains a location × features matrix of artificial neurons (ANs), with an array of ANs whose “feature-selectivity” is identical at each location. The RF-size of the ANs in different layers is determined by their connectivity, with ANs in deeper layers pooling over larger regions of space, similar to biological neurons in the primate ventral stream. The similarity of feature selectivity of ANs and biological neurons can be determined by comparing their responses across many images. Accordingly, for each image presented during the recordings, we computed the activation of artificial neurons (ANs) with convolutional RFs that overlapped with the RFs of the to-be-predicted recording sites (see Methods) (Figure 1H). For each VGG-16 layer separately, we then predicted neural activity using cross-validated sparse L1-regression from the activity of the ANs (as in Cadena et al. (46), Figure 1H-I). Note that the responses of the ANs in this case are simply regression predictors, which generate an output for each image that is then used to explain different kinds of neural activity (e.g. MU firing intensity or neuronal *γ* synchronization). Firing-rate intensity was computed separately for early (50-150ms) and late stimulus periods (200-600ms), by determining the instantaneous energy of the MU activity (see Methods). Note that in the manuscript, we primarily focus on the late rates, in which surround modulation was strongest (see further below).

V1 firing rates in the late period were relatively well predicted from the activation of VGG-16 ANs (Figure 1I), with maximum R values around 0.50 for the middle layers of the VGG-16 (3conv1 to 4conv2) (for early firing rates see Figure Extended Data 1A). This agrees with previous findings (Cadena et al. (46)) and with the convolutional RF-size in 3conv3, which is effectively 40×40 pixels (1dva; see Table 1). As expected from the fact that we centered the image around the neuronal RF, V1 firing rates were predicted by ANs around the center of the image, revealing small, circular RFs of around 1dva (Figure Extended Data 1B). For V1 LFP *γ*-synchronization, explained variance was generally much lower than for V1 firing rates (maximum R around 0.11) (Figure 1I) and as we will show later in the paper, much lower than can be obtained with alternative methods (Figure 3-5).

**Table 1:** Receptive Field sizes in the VGG-16 across layers. The abbreviation dva stands for degrees in visual angle. This was shown separately for monkey H and monkey A and I because of the differences in the number of pixels per degree.

|  |  |  |  |  |  |  |  |  |  |  |  |  |  |
| --- | --- | --- | --- | --- | --- | --- | --- | --- | --- | --- | --- | --- | --- |
| layer | 1c1 | 1c2 | 2c1 | 2c2 | 3c1 | 3c2 | 3c3 | 4c1 | 4c2 | 4c3 | 5c1 | 5c2 | 5c3 |
| pixels | 3 | 5 | 10 | 14 | 24 | 32 | 40 | 60 | 76 | 92 | 132 | 164 | 196 |
| dva (A & I) | 0.07 | 0.12 | 0.25 | 0.35 | 0.60 | 0.80 | 1.00 | 1.50 | 1.90 | 2.30 | 3.30 | 4.10 | 4.90 |
| dva (H) | 0.06 | 0.10 | 0.19 | 0.27 | 0.46 | 0.62 | 0.77 | 1.15 | 1.46 | 1.77 | 2.54 | 3.15 | 3.77 |

**Figure 3:**
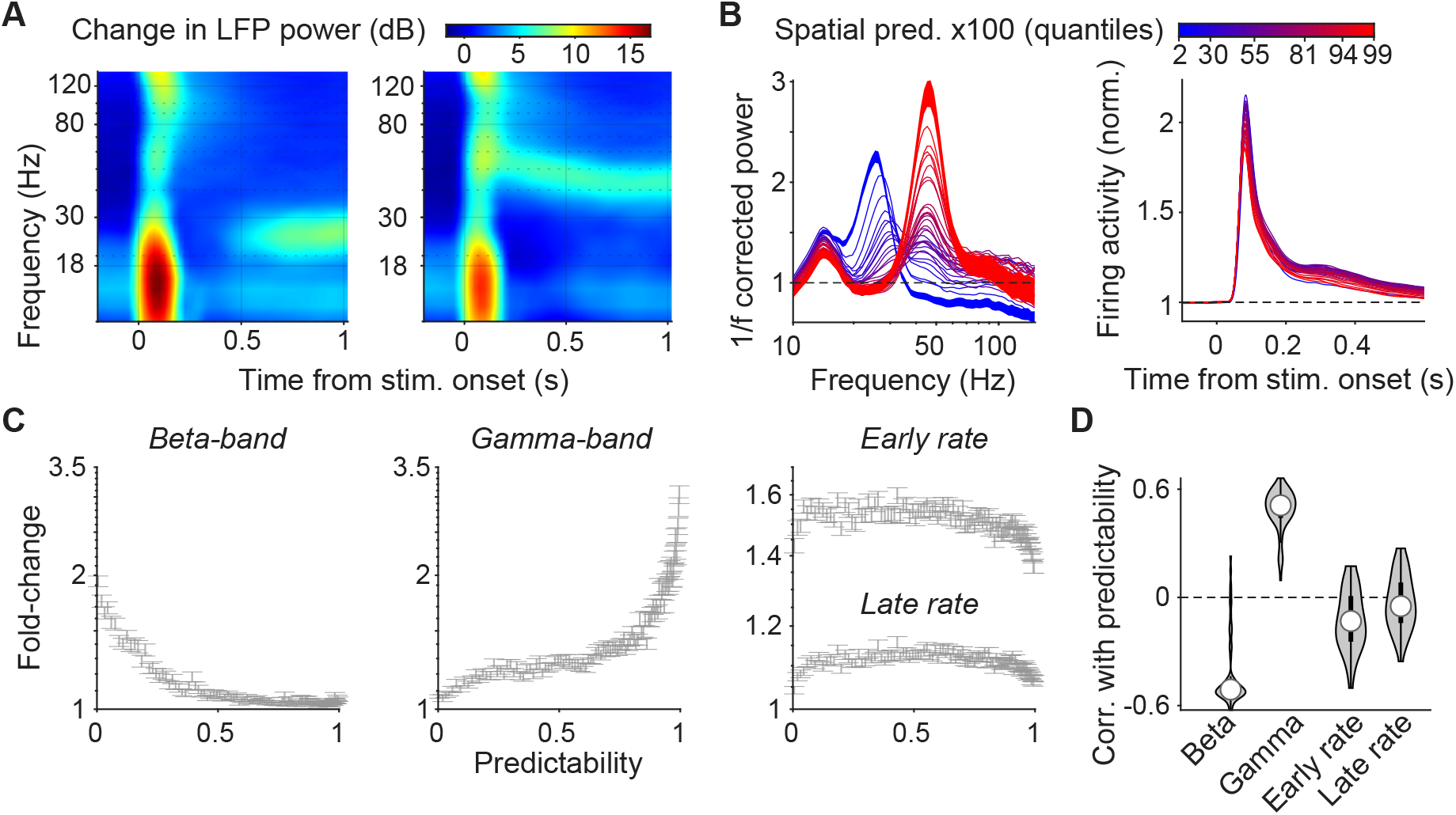
Distinct relationships of firing rates and neural synchronization with spatial predictability. **(A)** Average time-frequency representations for low (left) and high (right) spatial predictability, showing the late emergence of *β* (left) and early emergence of *γ*-synchronization (right). Spatial predictability was estimated as “structural predictability”, the pixel-by-pixel correlation between model-predicted and ground-truth image values (see Methods and main text). This quantifies the extent to which the precise pixel structure of a stimulus falling into the V1 receptive field can be predicted by the context. **(B)** Left: Average 1/f corrected LFP power spectra for monkey H. Spectra show distinct peaks in the *γ* and *β* frequency range. Blue-to-red colors indicate levels of spatial predictability. Black line indicates the pre-stimulus period. SEMs are shown for the lowest and highest quantile of spatial predictability. Right: Multi-unit firing rate for different levels of spatial predictability. **(C)** Left: Average (± SEM) of 1/f corrected *β*-peak amplitude depending on spatial predictability. Average was computed across all recording sites in the three animals (*N* = 72 sites). Middle: Same for *γ*. Right: Same for early (50-150 ms) and late firing rates (200-600 ms). **(D)** Average Pearson-r correlation across recording sites between spatial predictability and *γ, β* and rate. Correlations were computed for each channel separately, across all images presented across sessions. Correlations were significant for *β, γ* and early rates (all *P* < 0.001) but not for late rates (*P* = 0.11) (T-test). Absolute correlations were higher for *γ* and *β* than early and late rates (*P* < 0.001 for all comparisons).

### Structural stimulus predictability in natural images

A standard neural network for object recognition performs relatively well in predicting V1 firing rates from a natural image, but cannot account well for contextual modulation (47) and performs poorly on predicting V1 *γ*-synchronization, which is thought to reflect contextual relationships (37; 45). To study the contextual modulation of neural activity for natural images, we developed a new method to quantify spatial stimulus predictability of visual inputs to the RF in an self-supervised way. We trained a deep neural network in a self-supervised way to form stimulus predictions about a small region of the image based on the rest of the image, and improved these stimulus predictions through iterations, each time comparing the predicted image patch with the ground truth image patch. This can be understood as a form of predictive coding. The to-be-predicted part of the image will be referred to as the “RF image patch”. The input to the network was the embedding context, i.e. the image region surrounding the RF image-patch (224×224 pixels; approximately 5-6dva wide). Based on this contextual input, the network was trained to generate a predicted RF image-patch (the size of the generated output was set to 1dva, roughly corresponding to the size of the neuron’s RFs). The network was trained using a large number of natural images that were *not* presented during the recordings. Training was self-supervised, because only data from the image itself was used for training, without human labels. As a loss function for training, we quantified perceptual similarity between the input (ground-truth) RF image-patch and the predicted RF image-patch (see Methods). We then used this trained neural network to quantify spatial stimulus predictability for each of the images presented during the recordings, for each recording site separately (Figure 2; see Methods). For this, we used various measures of perceptual similarity to compare: (1) The *actual* visual input to the neurons, i.e. the RF image-patch. (2) The visual input that was *predicted* based on the rest of the image. We initially computed the pixel-by-pixel correlations (i.e. structural similarity) between the ground-truth and the predicted RF image-patch. We call the resulting measure the “structural predictability”, which reflects the extent to which the precise pixel structure of a stimulus falling in the V1 RF can be predicted by the context. Note that structural image similarity as quantified by pixel-by-pixel correlations is comparable to both the well-established perceptual-similarity measure SSIM (Figure Extended Data 2; see Methods); note that we omitted the dependence of SSIM on contrast and spatial frequency similarity to avoid an influence of low-level image factors.

We then examined the relationship between neural activity and the spatial predictability measure. LFP spectra were computed for 32 quantiles of structural stimulus predictability and differed markedly as a function of this spatial predictability measure (Figure 3A-B): For strong structural predictability, LFP power spectra showed a narrow-band peak in the *γ*-frequency range, which was on average several times larger than the pre-stimulus power (Figure 3A-B, Extended Data 3; for all time-frequency-representations see Figure Extended Data 4). Conversely, for weak structural predictability, LFP spectra did not show *γ*-peaks (Figure 3A-B). From the lowest to highest values of structural predictability, *γ*-synchronization increased monotonically and by approximately 300% (Figure 3A-C). This finding was consistent across the three monkeys (monkey H, I and A: Pearson’s r=0.94, 0.85, 0.87) and similar findings were made for MUA spiking activity in the gamma range (Figure Extended Data 5). In an additional analysis, we directly correlated LFP *γ*-power with structural predictability across all images, i.e. without using quantiles (see Methods). This revealed a clear positive correlation between LFP *γ*-synchronization and structural predictability (Figure 3D).

Unexpectedly, for weak structural predictability, LFPs showed a prominent peak in the (high) *β*-frequency band (18-30Hz) range (Figure 3A-B). This peak emerged only during the late phase of the stimulus period (>500ms) (Figure 3A; Figure Extended Data 4). *β*-synchronization with these stimulus characteristics has, to our knowledge, not been previously linked to sensory processing in area V1, but likely reflects late cortical feedback originating from higher brain regions like parietal and frontal cortex (48; 49; 34; 50). Opposite to *γ, β*-band activity showed a monotonic *decrease* as a function of structural predictability (Figure 3B-C). This was also found in multiunit (MU) activity, although *γ* fluctuations were overall more prominent in the MUA (Figure Extended Data 5). LFP *β*-peaks were detected in only 2/3 animals and were negatively related to structural predictability in both (Monkey H: Pearson’s r=-0.79; Monkey I: r=-0.54; note that in the third monkey, stimulus duration was only 600ms). For the rest of the paper we focus primarily on firing rates and gamma. For *β*, we report several of the main analyses in Figure Extended Data 6.

Early MU firing-rate intensity showed a weak negative correlation with structural predictability, and late firing rates were not significantly correlated (Figure 3B-D). The relationship between firing intensity and structural predictability was nonmonotonic, with maximal MU firing rates for intermediate structural predictability (Figure 3C). Thus, the relationship between firing rate and structural predictability was weak or nonsignificant, which appears to contradict the theory that V1 firing rates encode sensory prediction errors (1). However, we will show below that firing rates are determined by a different form of stimulus predictability, namely high-level stimulus predictability.

### Stimulus predictability and data compression

To further understand the significance of structural predictability for visual encoding, we directly related structural predictability to the data compression of the images. Images that contain predictable structure should be highly “compressible” because they have a high degree of redundant information. To quantify the relationship between compressibility and structural predictability, we used a large image database where we compressed each image and determined the number of bits per pixel in the compressed image thereby giving the data compression rate (see Methods). We then determined the average structural predictability across 16 locations in the image. This yielded a very strong correlation between the average structural predictability and compressibility (Figure 4A-B), showing that images with high structural predictability can also be efficiently encoded.

**Figure 4:**
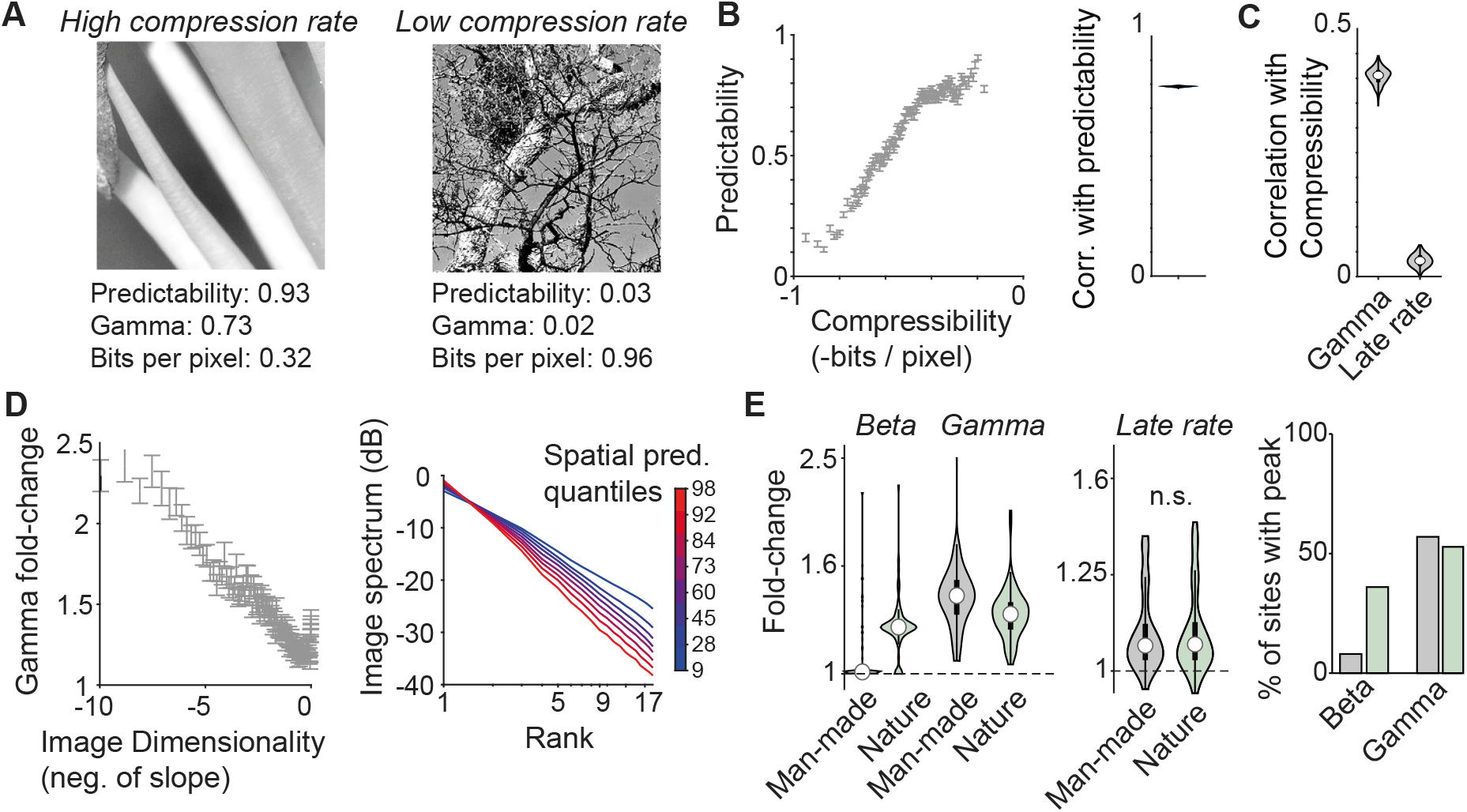
Relationships of firing rates and neural synchronization to compressibility, dimensionality and natural image category. **(A)** Two examples of images that have a low and high compression rate, spatial stimulus predictability and *γ* values, respectively. Compression rate was measured as the number of bits per pixel for image compression. **(B)** Left: Compressibility (i.e. negative of compression rate) vs. average spatial predictability for natural images, ordered by quantiles. Right: Correlation of compressibility with spatial predictability across images. **(C)** Correlation between compressibility and *γ* and firing intensity, across images. Correlations were computed per channel. **(D)** Left: Images with low dimensionality had strong *γ* synchronization (R across quantiles = −0.9, *P* < 0.001). Dimensionality was determined by first taking the two-dimensional fast Fourier transform of the RF image-patch, taking the rotational average, and ranking the spectral components by magnitude. Dimensionality was then defined as the slope of the resulting spectrum. An image with lower dimensionality can be represented by fewer spatial frequency components. Right: Average magnitude of spectral image components for different quantiles of spatial predictability (Pearson’s R between dimensionality and predictability across quantiles = −0.91, *P* < 0.001). **(E)** Fold-changes in neural activity for images with man-made content (e.g. buildings) in the RF and images with nature content (e.g. trees, bushes) in the RF. Comparison was significant for *β* (*P* < 0.001) and *γ* (*P* < 0.001), but not for early and late rates (*P* = 0.5 and *P* = 0.9, T-Test). Right: Percentage of sites with a *β* or a *γ* peak, across all randomly selected images.

Based on Figure 3, this would entail that *γ* but not firing rates correlate strongly with compressibility. To investigate this, we computed the compressibility of the 3 × 3 image patches around each RF and correlated this with neural activity. We found that *γ* showed a much stronger correlation with compressibility than firing rates (Figure 4C, Extended Data 7).

### Predictability, dimensionality and natural image statistics

Next, we wondered how neural activity relates to the dimensionality of the visual input. We expected that when the RF input can be well predicted by the surrounding context (e.g. a grating or a homogeneous surface), the image can be represented by relatively few spectral components. This can be captured by the concept of “dimensionality”. To quantify dimensionality, we took the rotational average of the two-dimensional Fourier transform of the RF image-patch, we sorted the spectral components by their magnitude and defined dimensionality as the slope of the spectrum. For image patches with higher structural predictability, we observed lower dimensionality (Figure Extended Data 8). Likewise, *γ*-synchronization was maximal for images with low dimensionality and showed a monotonic decrease towards high dimensionality (Figure 4D). This shows an interesting parallel between dimensionality at the neuronal and image level: low-dimensional stimuli are mapped onto low-dimensional neural states. More specifically, when the image has a low dimensionality, in the sense that it can be reconstructed by fewer spectral components, neural activity is also dominated by energy in a relatively narrow frequency band (*γ*) as reflected by population-wide synchronized firing.

We then asked if neural activity could distinguish between different image categories (9). We observed that images with predictable structure often contain man-made objects, whereas unpredictable structure is common for pictures of nature (Figures Extended Data 10 and Extended Data 9). Furthermore, predictable structure is associated with the presence of object boundaries covering the RFs. Firing rate intensity did not differ between nature and man-made picture categories, and did not depend on the presence of an object boundary (Figure 4E and Figure Extended Data 11). Yet, *β*-synchronization was relatively strong for pictures of nature, whereas *γ*-synchronization was relatively strong for pictures with man-made content and an object boundary in the RFs (Figure 4E, Figure Extended Data 11). Consistent with this, simple artificial stimuli like gratings, straight and curved bars, and edges of filled contours also generated strong *γ* (Figure Extended Data 12, Extended Data 13, Extended Data 14, Extended Data 15), although some natural stimuli generated stronger *γ* than gratings recorded in the same session (Figure Extended Data 12). As a consequence of the near uniform distribution of structural predictability in natural images (Figure Extended Data 10), there was substantial variation in *γ* and *β* across images: *γ*-peaks were detected only for about 50% of sites, and *β*-peaks were found almost exclusively for the nature category (Figure 4E).

We further analyzed the influence of color, which can influence both *γ*-synchronization and firing rates (51; 45; 52; 53), with highly spatially predictable, homogeneous surfaces inducing strong *γ* synchronization and sparse firing (45). In separate sessions, we therefore presented images both in grayscale and in their original color. Across images, *γ* and *β*-synchronization both showed a strong correlation between grayscale and color images (Pearson’s r=0.81 and 0.7, respectively, *P* < 0.001 for both). Thus, the relationship between *β*/*γ* and structural predictability at the grayscale level explains substantial variance in *β*/*γ* for natural color images.

### Interaction of stimulus predictability with stimulus drive

We wondered how other image parameters influenced synchronization and firing rates and interacted with spatial predictability. For instance, luminance contrast shows substantial variability in natural images (see Figure Extended Data 10C) and can change the contextual modulation of sensory responses (54; 10; 55). For each recording site, we computed the luminance contrast of its corresponding RF image-patch (as RMS contrast; see Methods). Firing rates showed a strong increase as a function of luminance contrast (Figure 5A-C). Furthermore, luminance contrast correlated positively with *γ*-synchronization (Figure 5A-C; for *β* see Figure Extended Data 6), and the *γ* peak frequency showed a moderate increase with contrast (Figure 5A), consistent with previous work using grating stimuli (56; 57; 58).

**Figure 5:**
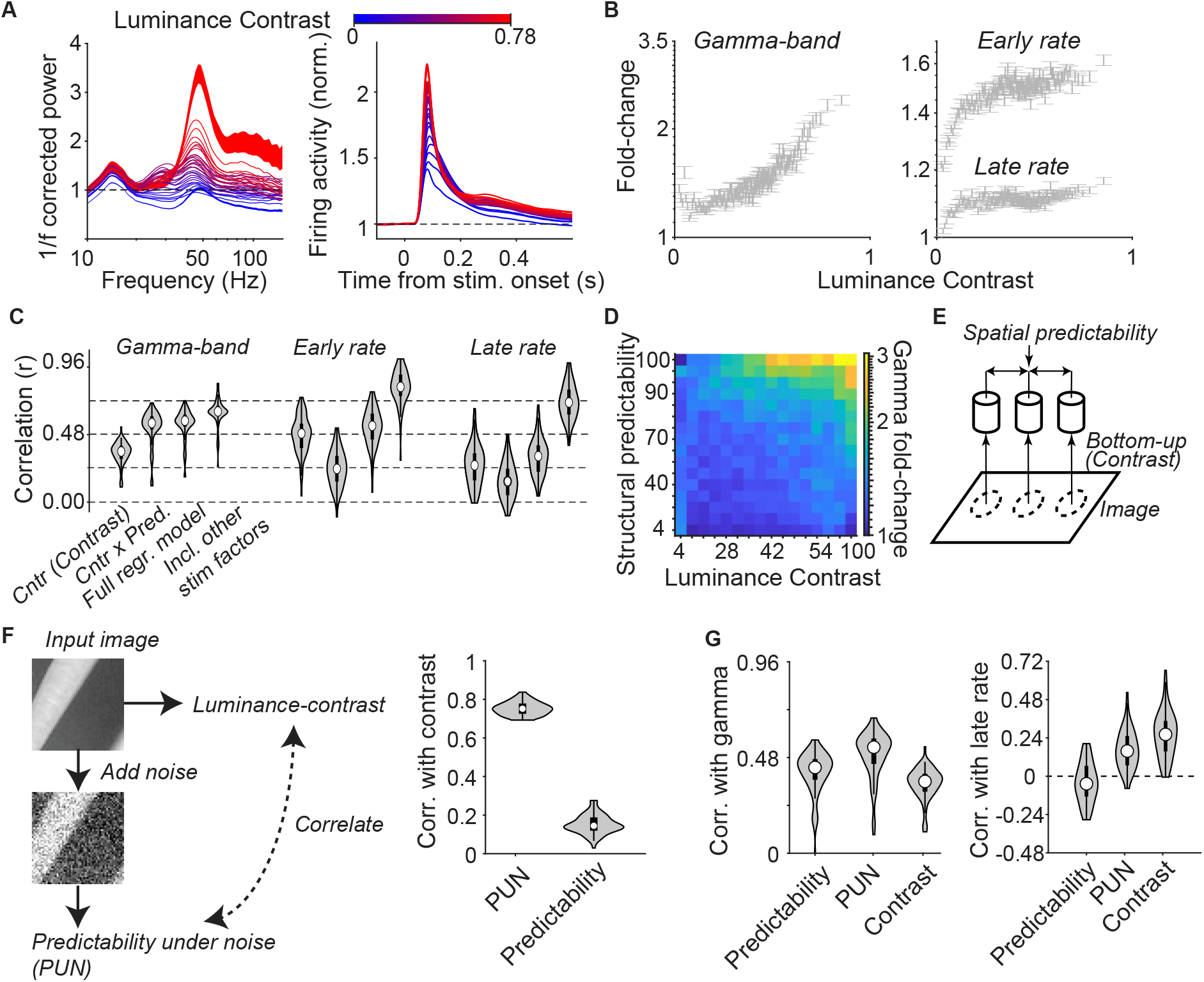
Dependence of firing rates and synchronization on luminance contrast. **A)** Left: Average 1/f-corrected LFP power spectra, with SEMs shown for the highest and lowest level of luminance contrast, for monkey H. Right: Multi-unit firing rate for different levels of luminance contrast. **(B)** Left: *γ*-peak amplitude for different levels of luminance contrast. Right: Same for early MU firing rates (50-150ms) and late MU firing rates (200-600ms). **(C)** Average correlation across sites of *γ* and firing rate with image factors. Left-to-right: Luminance contrast (Cntr); the product of contrast and stimulus predictability (Cntr x Pred.); predictability; contrast, predictability × contrast interaction, (Full regr. model); and a model with additional low-level features (Incl. other stim factors), namely spatial frequency, luminance and orientation (see Methods). The correlation was computed for each recording site separately across all images presented across sessions. All correlations were significantly different from zero (*P* < 0.001, paired T-test). For *γ*, the difference between (Cntr x Pred.) and (Full regr. model) was significant (*P* < 0.001), but the difference between (Full regr. model) and (Incl. other stim factors) was not (*P* > 0.05). For early rates, all comparisons were significant (*P* < 0.05). For late rates, all comparisons except for (Full regr. model) vs. (Cntr x Pred.) were significant at *P* < 0.05. **(D)** Heatmap of *γ* fold-changes for different quantiles of luminance-contrast and spatial predictability. **(E)** Illustration of interaction between predictability and bottom-up inputs. Firing rates and synchronization can be modified both by bottom-up inputs and image predictability. The influence of spatial predictability is mediated by horizontal, recurrent interactions or top-down inputs. **(F)** Derivation of the PUN measure (Predictability under noise, left). We added Gaussian noise to each image and then computed the structural predictability for the noise-corrupted image, yielding the PUN measure. Gaussian noise had a mean of 128 and a standard deviation of 42 pixel values. Pixel values were clipped at [0, 255]. This was motivated by the idea that V1 does not receive a noise-free representation of the original image, but an image represented by sparse and variable neuronal inputs. The PUN measure was strongly correlated to the original luminance-contrast in the center RF (right), whereas luminance-contrast was weakly correlated to structural predictability computed on the original image. **(G)** Left: PUN correlated more strongly with *γ* synchronization than predictability and luminance-contrast (*P* < 0.001 for both, paired T-test). Right: However, for late rates, correlations with PUN were weaker than with luminance-contrast (*P* < 0.001).

Because *γ*-synchronization was positively correlated both with structural predictability and luminance contrast, we wondered how these two variables interacted. We have previously suggested that stimulus predictability and luminance contrast interact in a multiplicative way, based on the idea that inputs to V1 neurons become more predictable / redundant under conditions of high signal-to-noise ratio (45). To investigate this, we fitted a multiple linear regression model, using structural predictability (Figure 3), luminance contrast and their interaction as predictors. This full regression model explained substantially more variance in *γ*-synchronization compared to luminance contrast or predictability alone (Figure 5C). There was a significant interaction between luminance contrast and structural predictability, with a higher T-statistic than for the individual variables (mean T-statistic contrast −1.42± 0.89, *p* > 0.05, T-test; predictability −0.84±0.78, *p* > 0.05; predictability × contrast 4.2 ± 1.2, *p* < 0.001). The interaction term of luminance contrast and structural predictability explained almost as much variance as the full regression model (Figure 5C). This multiplicative effect is consistent with the absence of *γ*-synchronization for both small, low-contrast stimuli, and highcontrast stimuli with low structural predictability (Figure 5D-E, Figure 7).

**Figure 6:**
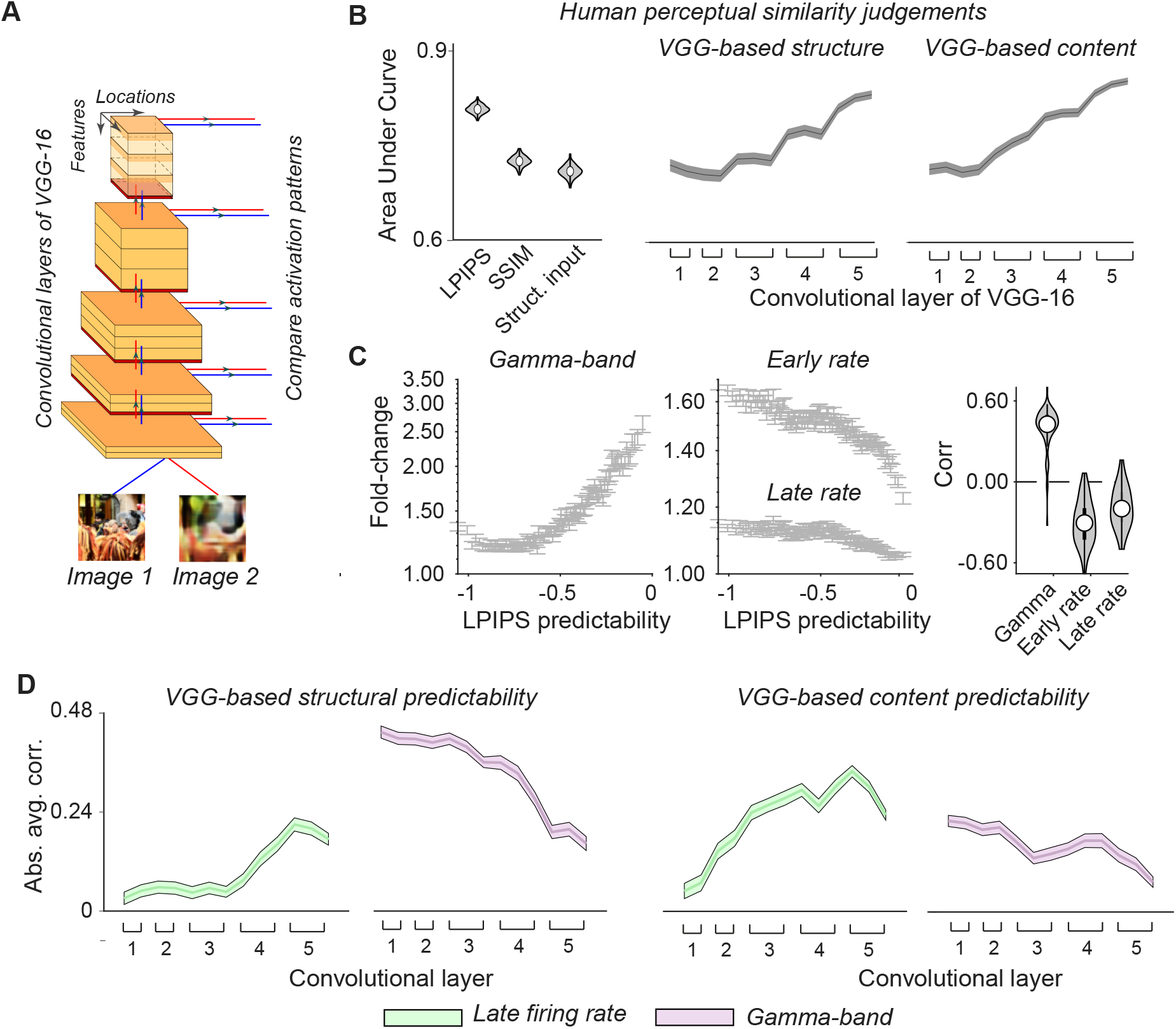
Modulation of V1 activity by the perceptual similarity measure LPIPS and low vs. high-level stimulus predictability. **(A)** VGG-16 network used to define low- and high-level stimulus similarity and predictability. Responses of VGG-16 artificial units (ANs) in different layers were computed both for two images at a time (e.g. the input (ground-truth) and predicted image-patch). For each VGG-16 layer, we computed two similarity measures: Content similarity (based on Euclidean distance) and VGG-based structural similarity. VGG-based structural similarity was computed in analogy to pixel-by-pixel correlations, by computing the Pearson correlation across locations for each AN separately and then averaging these across ANs. **(B)**: Left: Average AUC value for (1) LPIPS, (2) SSIM, (3) Structural correlations, as used for Figure 2 and 3. LPIPS (Learned Perceptual Image Patch Similarity) is a perceptual similarity measure based on deep neural networks for object recognition (60). LPIPS showed superior performance compared to the other measures (*P* < 0.001, Paired T-test). Right: Structural and content similarity vs. human perceptual similarity. AUC increases significantly with layer depth for both structure and content (R = 0.93 and R = 0.98, *P* < 0.001 for both), indicating that activation of units in deepest layers of the network explained human perceptual similarity judgments the best. **(C)** Correlations of firing rates and neural synchronization with spatial stimulus predictability defined as LPIPS. Left: Same as Figure 3C, but now with predictability defined as LPIPS. Right: Correlations across all images, averaged across recording sites; correlations were significant for all variables (*P* < 0.001). **(D)** Correlation of late firing rate (200-600ms, green) and peak *γ*-power (red) with VGG-based content and structural predictability across VGG-16 layers. Note that we first computed average correlations and show here the absolute average value of these correlations; correlations were positive for *γ* and negative for firing rates. Late firing rates showed a significant increase in absolute correlation across VGG-16 layers, both for VGG-based structural and content predictability (structure: *R* = 0.85; content: *R* = 0.81, *P* < 0.001 for both). By contrast, *γ* showed a significant decrease (structure: *R* = 0.9; content: *R* = 0.84, *P* < 0.001). The average correlation (across layers) for VGG-based structural predictability was significantly higher than VGG-based content predictability for *γ* (*P* < 0.001, Paired T-test). For firing rates, the average correlation with VGG-based content predictability was higher than for VGG-based structural predictability (*P* < 0.001, Paired T-test).

We wondered whether luminance contrast might, like structural predictability, reflect the presence of a high amount of redundant information in the visual inputs to V1. The rationale behind this is that V1 does not have access to the image itself, but to a potentially noisy representation of the image, encoded by sparse and variable LGN inputs. To investigate this, we added Gaussian white noise to the original images and quantified the resulting predictability as described above for each image (Figure 5F). This yielded a structural predictability value for the noisy images, i.e. the extent to which a noisy RF input could be predicted by the noisy image context (predictability under noise - PUN). The original luminance contrast correlated very strongly with PUN (Figure 5F). Furthermore, PUN correlated more strongly with *γ* synchronization than the structural predictability computed over the original (i.e. noise-free) image (Figure 5G). For firing rates however, correlations with PUN were weaker than with the original luminance-contrast (Figure 5G). This finding suggests that it is the structural predictability at the level of actual neuronal input (a noisy representation of the image) that determines *γ*-synchronization, instead of the structural predictability at the level of the image, consistent with the multiplicative effect of luminance-contrast and stimulus predictability.

We further examined whether other stimulus factors explained further variance (see Methods). We first examined the correlation of *γ*-synchronization with image focus (i.e. as opposed to blur), which has previously been shown to correlate with gamma (59). Image focus was strongly correlated with luminance contrast and showed weaker correlations with *γ*-synchronization than luminance contrast (Figure Extended Data 16). Next, we included additional regression predictors namely spatial frequency, stimulus orientation and luminance. These variables were computed both for the 1dva center patch and the 224×224 image region. Including these additional regression predictors only slightly increased the explained variance for *γ*-synchronization (Figure 5C). Structural predictability did not interact with spatial frequency in a multiplicative manner (Figure Extended Data 17). In contrast to *γ*, including these low-level image factors had a very strong effect on explained variance for firing rates, consistent with the Gabor-like RF properties of V1 neurons (Figure 5C). The distinct influence of low-level stimulus factors on *γ* and firing rates matches with their respective correlations to the activity in early layers of the VGG-16 (Figure 1).

### Perceptual similarity and low vs. high-level predictability

In Figure 3, we showed that late firing rates were not significantly correlated with structural predictability, i.e. the extent to which the precise stimulus structure into the V1 RF was predicted by the context. The precise pixel structure of an image has been widely used to determine the perceptual similarity between two images (SSIM). However, recently, new methods for perceptual similarity have been developed which outperform these traditional perceptual similarity measures. These methods compare images not only in terms of low-level, but also high-level features (60). Specifically, LPIPS (Learned Perceptual Image Patch Similarity) compares two images based on the similarity of their representations across all layers of a CNNs for object recognition (Figure 6A). We first replicated the finding that LPIPS correlated better with human perceptual similarity than SSIM or pixel-by-pixel correlations (structural similarity) (Figure 6B, Extended Data 18). The difference between LPIPS and SSIM can be explained by the observation that perceptual similarity judgments correlated most strongly to patterns of activation in the *deep* layers of the VGG-16 (Figure 6B). This held true for two standard types of image similarity, VGG-based content and structural similarity (Figure 6B). VGG-based content is based on the Euclidean distance between activation patterns in the VGG-16 (61). VGG-based structure is based on the Pearson correlation between activation patterns across locations, in analogy to the pixel-by-pixel structural similarity or SSIM (thus, this weights the spatial correlation pattern more compared to content).

We then applied these VGG-based similarity measures to quantify spatial predictability. Neurons can in principle encode stimulus predictions or prediction errors of varying levels of complexity, based on either low-level structure (e.g. pixel values) or high-level features that can provide information about the presence of different objects. We thus wondered if firing rates would be modulated by a higher-order form of predictability. To investigate this, we used the LPIPS measure to compare the ground-truth and predicted image (using the algorithm detailed in Figure 2), yielding “LPIPS-predictability”. In contrast to (pixel-by-pixel) structural predictability, LPIPS-predictability also captures how well higher-level features of the stimulus in the RF are predicted by context. In contrast to structural predictability (Figure 3), we found that V1 rates in the late response phases were negatively correlated and monotonically related to LPIPS-predictability (Figure 6C). Further, correlations between V1 *γ* and LPIPS were comparable to correlations between V1 *γ* and structural predictability (Figure 6C). This likely reflects the fact that LPIPS is based on both low-level structure and high-level features (see further below).

The observation that LPIPS-predictability (Figure 6C) but not structural predictability (Figure 3) correlated with rates, suggests that firing rates might be sensitive to a higher-level form of stimulus predictability, rather than the predictability at the level of pixel-by-pixel correlations. To investigate this, we defined layer-specific content and VGG-based structural predictability (using the VGG-based content and structural similarity) (Figure 6D). For both stimulus predictability measures, we found that rates were negatively related to predictability, and that the magnitude of the correlation between stimulus predictability and V1 firing rates became stronger towards the deeper layers of the VGG-16 (Figure 6D; showing the magnitude of the average correlation). Further, V1 firing rates were better explained by layer-specific content predictability than VGG-based structural predictability (Figure 6D, Figure Extended Data 19)). Thus, high-level content predictability was the most reliable variable explaining V1 firing rates.

We observed the opposite pattern for V1 *γ*-synchronization (for *β* see Figure Extended Data 6): V1 *γ* was strongly and positively correlated with VGG-based structural and content predictability in the early layers of the VGG-16 (Figure 6D), but weakly correlated with content and VGG-based structural predictability in the deeper layers of the VGG-16. Accordingly, the correlation with VGG-based predictability decreased across layers (Figure 6D). Furthermore, correlations with layer-specific content predictability were substantially weaker than with VGG-based structural predictability (Figure 6D). Including VGG-based structural and content predictability from deeper layers did not improve the prediction of V1 *γ*synchronization, in contrast to V1 firing rates (Figure Extended Data 19).

These findings indicate that *γ*-synchronization and firing intensity reflect distinct aspects of stimulus predictability: Firing rates decrease predictability of high-level features, whereas *γ* increases with low-level predictability. Whereas high-level features predict human perceptual similarity judgeents well, the opposite holds true for image compression: Compressibility correlated best with the activity in early, rather than deep layers of the VGG-16 (Figure Extended Data 7).

### Experimental dissociation of firing rates and *γ*-synchronization

We designed a further experiment to dissociate firing rates from *γ*-synchronization. Noise textures (brown, white, pink) were presented in two conditions: Center-surround match (same noise in RF and surround) and mismatch (different noise type in RF and surround). This paradigm can be compared to the classic figure-ground paradigm albeit it with a figure of the size of a RF (62). Because these stimuli had low structural predictability, we did not expect to find any *γ*-synchronization. Indeed LFP spectra did not show *γ* peaks for any of the conditions (Figure 7A). However, these stimuli differed in terms of higher-level content predictability, since ANs in deeper layers of the VGG-16 have texture selectivity. Accordingly, firing rates were substantially higher for texture mismatches than homogeneous textures (Figure 7A). These findings further support the conclusion that firing rates and synchronization properties are modulated by distinct aspects of spatial predictability.

**Figure 7:**
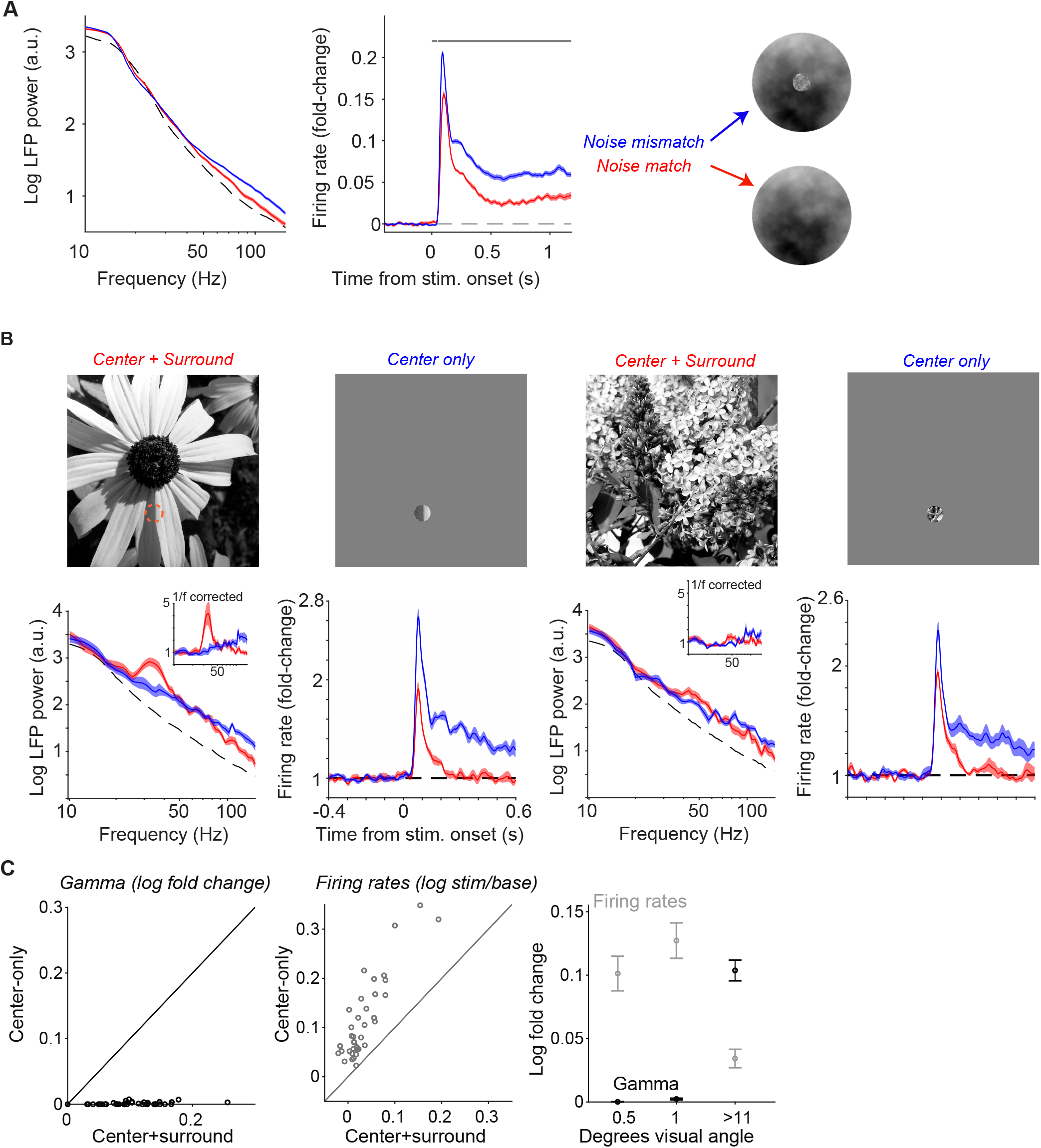
**(A)** Center-surround mismatch paradigm with noise stimuli and center-only vs full stimuli. Stimuli had white, pink or brown noise in the 1dva center, and white, pink or brown noise in the surround. Stimulus size was 6dva. Stimuli were centered on the recording site’s RF. Only recording sites within 0.25 dva of the stimuli center were included in the analysis. Left: LFP power-spectra for noise mismatch and noise match stimuli. Note that the broad-band increase in LFP power at high frequencies is typical for spike bleed-in (41). The dashed line represents the baseline pre-stimulus period. Right: Normalized MU firing-rates for noise match and mismatch stimuli. Firing rates were significantly higher for noise-mismatch stimuli (gray bar: *P* < 0.05, T-test, N=24 recording sites). **(B)** Left: Example of surround suppression for an image that shows clear *γ*-synchronization. Right: Example of surround suppression for an image that shows no clear peak in the *γ*-range. **(C)** Left: Comparison of *γ*-synchronization (log_10_ fold-change) between the center+surround (full image) vs. center-only (0.5 and 1dva pooled). Middle: Surround suppression for the late firing-intensity, and showing log_10_(*stim/base*). Suppression for early firing-intensity was significantly weaker than for late firing-intensity (paired T-test, *P* < 0.001). Right: Averages and standard errors of the mean for the different stimulus sizes.

### Stimulus-size dependence and prediction of surround modulation

Next, we investigated whether the surround modulation of neural activity, i.e. the difference between small and large stimuli, was explained by stimulus predictability. To further investigate the dependence of neural activity on context, we recorded separate sessions in which images were presented either in full (“center+surround”) or within a small (0.5-1dva) aperture (“center-only”) centered on the RFs of the recorded neurons. This served two purposes: First, it allowed us to dissociate the dependence on context from local stimulus properties, keeping the local properties constant while exclusively manipulating the surround. Second, it allowed us to investigate if surround modulation itself was determined by stimulus predictability. The state-of-the-art measure to quantify center-surround homogeneity for natural images is based on the first- and second-order statistics in natural images. Specifically, responses of Gabor RF filters (similar to simple cells) show correlated variance across natural images (63). Based on this observation, Coen-Cagli et al. (13) derived a Bayesian measure of homogeneity (P(homogeneity), and achieved improved prediction of firing-rate surround-modulation compared to previous models (in anesthesized monkey V1) (13). In contrast to the stimulus predictability measures developed here, P(homogeneity) is a bimodally distributed probability function that does not distinguish different levels of stimulus predictability in a continuous way like LPIPS. Furthermore, because P(homogeneity) is based on Gabor-like RFs, it may not capture aspects of highlevel stimulus predictability.

We found that firing rates were consistently reduced in the center+surround condition (Figure 7B-C), consistent with previous work (14; 13). Surround suppression was stronger for late than early firing rates (pairwise T-test, *P* < 0.001, Early: 0.043+/-0.009, Late: 0.08+/-0.008) consistent with dependence of surround suppression on horizontal and top-down feedback (17). In contrast to firing rates, *γ*-synchronization was strongly reduced in the center-only condition (Figure 7C). Thus, the emergence of *γ* required that there was a RF “surround”, i.e. that the natural image extended beyond 1dva with a predictable context (Figure 7C). This indicates that low-level RF properties alone were not sufficient to induce *γ*-synchronization.

Surround suppression was sometimes associated with strong *γ*-synchronization and in other cases occurred in the absence of a clear *γ* peak (Figure 7B). To further understand how surround modulation of rates and *γ* was determined by contextual relationships, we correlated stimulus predictability (computed for the full image) with surround suppression, for each recording site separately (Figure 8A). LPIPS-predictability correlated more strongly with surround suppression of rates than P(homogeneity) and the low-level structural predictability (Figure 8B-C). Although rate surround suppression was already found for low stimulus predictability, surround suppression increased monotonically across different levels of LPIPS-predictability (Figure 8C). By contrast, it did not show a significant trend across different levels of P(homogeneity), nor with structural predictability (Figure 8C). This is expected, given that LPIPS-predictability is a continuous and more uniformly distributed variable, whereas P(homogeneity) is a bimodally distributed measure (13).

**Figure 8:**
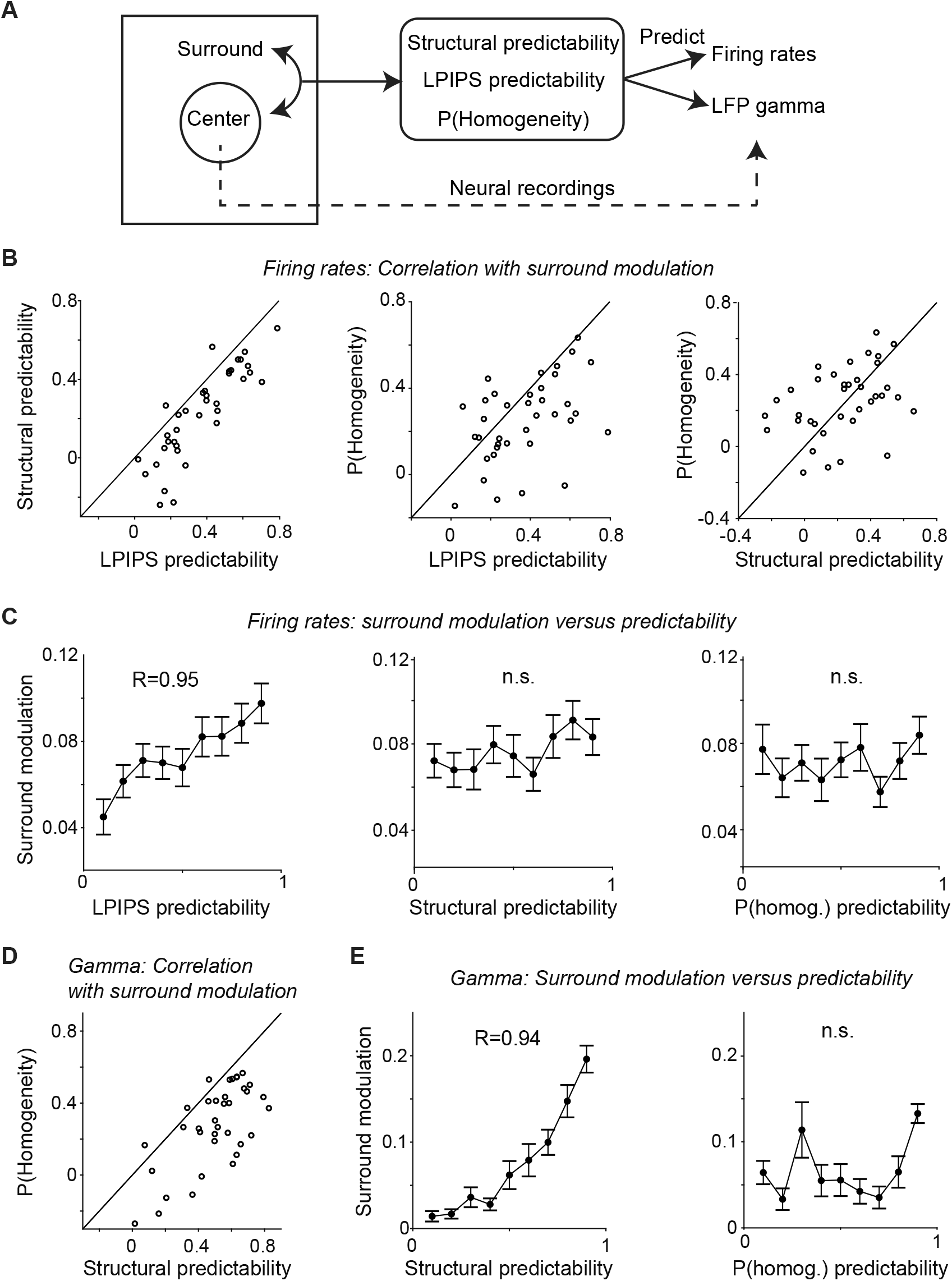
Stimulus predictability and surround modulation of firing rates and *γ*-synchronization. **(A)** Stimuli were presented in the masked (0.5 and 1dva) or full configuration, as shown in Figure 7. For each image, we computed structural and LPIPS predictability based on the 4 × 4 dva patch around the center. In addition, we computed P(homogeneity), the center-surround homogeneity measure of (13), a bimodally distributed, probabilistic measure of center-surround homogeneity based on energy correlations between Gabor-like simple-cell responses (13). These measures were then correlated with the surround modulation of firing rates and *γ*-synchronization. **(B)** Left: Spearman correlation between surround modulation and LPIPS-predictability vs. Spearman correlation between surround modulation and structural predictability. Each circle corresponds to a channel for which the stimulus was centered on the channel’s RF. LPIPS-predictability correlated more strongly with surround modulation than structural similarity (*P* < 0.001, pairwise T-test). Middle: Same as B-left, but now on the y-axis the correlation between surround modulation and P(homogeneity). LPIPS-predictability correlated more strongly with surround suppression than P(homogeneity) (*P* < 0.01, pairwise T-test). Right: Same as B-left, but now contrasting structural predictability and center-surround homogeneity. Difference was non-significant. **(C)** Left: Average surround modulation (center-only minus center+surround) as a function of different levels of LPIPS predictability. Correlation across levels was significant (*R* = 0.95. *P* < 0.001). Middle: Same as left, but now for structural predictability. Right: Same as middle, but now for P(homogeneity). Correlation was not significant (*P* > 0.59). **(D)** Spearman correlation between surround modulation of *γ*-synchronization and structural predictability vs. Spearman correlation between surround modulation and structural predictability. Correlation was stronger for Structural Predictability than P(homogeneity) (*P* < 0.001, pairwise T-test). **(E)** Left: As C-middle, but now for *γ*. Correlation across levels was significant (*R* = 0.94. *P* < 0.001). Right: As C-right, but now for *γ*. Correlation was not significant (*P* = 0.49).

Nonetheless, LPIPS-predictability showed a significant correlation with P(homogeneity) of *ρ* = 0.5 (*P* < 0.001) for our images. To further understand the redundancy between these factors, we performed a multivariate regression analysis, and found that LPIPS and not P(homogeneity) was the dominant regression predictor of surround modulation (LPIPS-predictability: *T* = − 1.60 + */* − 0.17, P(homogeneity): *T* = − 0.27 + */*− 0.13; note that we used log(P(homogeneity)) as it showed a stronger linear correlation with LPIPS).

In contrast to firing rates, surround modulation of *γ* increased monotonically with structural predictability, consistent with the findings above (Figure 8D-E). Furthermore, *γ*-synchronization showed a weaker relationship with P(homogeneity) than with structural similarity (Figure 8D). Thus, the stimulus predictability measures developed here allow for a continuous regression prediction of the surround modulation of firing rates and *γ* synchronization, which are modulated by distinct types of stimulus predictability (see also Figure 6).

## Discussion

We investigated how two distinct features of V1 activity, firing rates and neuronal synchronization, are modulated by predictive spatial context. Recordings were made in awake monkeys, which is important because contextual modulation depends on brain state (64; 16).

According to predictive and efficient coding models, V1 neurons should fire sparsely when they receive bottom-up inputs that are predicted from the spatial context (1; 65). To measure this predictability in natural images, we first trained a deep neural network to predict the image in the neurons’ RF region from the context, the non-RF region (Figure 2). We then derived a continuous measure of predictability by comparing the predicted image with the ground-truth image using different perceptual similarity functions. Initially, we defined the structural predictability, by quantifying the loss at the pixel-by-pixel level using a classic measure of image similarity, i.e. structural similarity (closely related to perceptual similarity SSIM). The structural predictability indicates to what extent the precise spatial structure of a stimulus falling into a V1 RF can be predicted by the context, and was strongly correlated with the compressibility of the natural image and its dimensionality. Surprisingly, V1 firing intensity was not significantly correlated with structural predictability (Figure 3), which appears inconsistent with efficient coding theories.

Notably, *perceptual* similarity is mainly determined by higher-level features of natural images, not by the precise pixel structure (Figure 6, Zhang et al. (60)). Likewise, predictability can be defined in terms of both lower level precise pixel structure and higher-level stimulus features, which can be quantified using CNNs (Figure 6). For example, the fur of an animal might be weakly predictable in terms of pixel-by-pixel correlations, but could be highly predictable for (artificial) neurons with complex feature selectivity thereby providing key information to distinguish between animals. When higher-level features of stimuli falling into the V1 RF were predicted by the context, V1 neurons showed a strong reduction in firing rates. This higher-level feature estimation allowed us to achieve improved (regression) prediction of surround suppression as compared to state-of-the-art predictors (Figure 8) (13). Together, these findings suggest that predictive and efficient coding of the image by V1 firing rates applies specifically to high-, but not low-level stimulus predictability. It is unclear how to reconcile this discovery with standard predictive coding models, in which V1 neurons are hypothesized to compute the difference between bottom-up inputs into V1 and top-down predictions about those inputs (1). This also raises the possibility that sensory prediction-errors do not propagate exclusively in the *feedforward* (FF) direction, and suggests that stimulus prediction errors may also flow in the FB (feedback) direction (see also Issa et al. (22); Heeger (20); Schwiedrzik and Freiwald (21)).

FF communication is thought to depend on synchronization of output signals (66; 67), which is promoted by V1 *γ*-oscillations (23). Recent work suggests that directed functional connectivity in the FF and FB direction is primarily signaled in the *γ* and *β* frequency-range, respectively (48; 35; 34) (but see Schneider et al. (36); Schomburg et al. (68)). Some groups have therefore hypothesized that *γ*and *β*-synchronization sub-serve the transmission of FF prediction errors and FB predictions, respectively (31; 69; 32; 33). Our findings, however, seem incompatible with this hypothesis: (1) *γ*-synchronization emerged with high rather than low structural predictability (i.e. low stimulus prediction error) (Figure (Figure 3, 6). (2) *β*-synchronization, which emerged in the late phase of the trial and likely reflects top-down FB (70; 71; 48), was induced by weakly predictable image patches with primarily nature content (Figure 3, 4). Thus, we found that firing rates and *γ*-synchronization carried complementary and independent information for natural scenes: Firing rates decreased with high-level stimulus predictability, whereas *γ* increased with low-level stimulus predictability. Furthermore, firing rates were well explained by the activity of artificial neurons in standard feedforward CNNs for object recognition, yet *γ* was not.

Our results on *γ* rather support the hypothesis that *γ*-synchronization reflects center-surround predictability, and the mutual predictability of visual inputs at different RF locations (“synchronization through mutual predictability”; STMP) (37). Our results further demonstrate that this principle pertains specifically to low-level structural predictability. Related to the STMP principle are recent computational models showing that *γ* synchronization can emerge in E/I networks performing probabilistic inference based on a generative model of natural image statistics, or efficiently encoding afferent inputs (72; 73). Because low-level image representations should correspond to inputs into V1, we conceptualize *γ*-synchronization as a signal that reflects the extent to which the precise structure of a stimulus in the V1 RF can be predicted from the context, and thereby provides redundant and compressible information (37). Consequently, the image regions providing redundant information engage in *γ*-synchrony (45), whereas image regions that provide more unique information exhibit asynchronous, irregular spiking. Importantly, we obtained these results on stimulus predictability using learned image statistics rather than specifying a restricted set of human-defined features as in previous studies on contextual modulation (74; 45; 23; 75; 37; 39; 76), and with a large and carefully controlled natural image set. This is critical because contextual stimulus predictions should follow the statistics of the natural environment. The concept of STMP (37) explains many of the basic feature-selectivities of V1 *γ*,e.g. checkerboards (77), stationary gratings (78), straight and curved stationary bar stimuli and edges of filled contours (Figure Extended Data 13,Extended Data 14,Extended Data 15), and homogeneous colored surfaces (45; 51; 79). As an example, we have previously shown that a large red colored surface induces *γ* synchronization everywhere, however a small green blob of color induces a near-complete loss of *γ* synchronization at the location of the green blob (45). Our findings suggest that the multiplicative dependence on luminance-contrast is also consistent with the principle of stimulus predictability and redundancy. Importantly, the emergence of *γ* requires a non-linear computation over center and surround inputs, because a linear combination of CNN filter responses only weakly predicted *γ* (Figure 1I, Figure 6 in Uran et al. (80)). In general, these findings agree with the original insight of Gray et al. (23) that *γ*-synchronization reflects feature relationships. Furthermore, the emergence of *γ*-synchronization likely implies a grouping according to local perceptual Gestalt principles (81). However, the inverse is likely not the case: not all forms of local Gestalt principles will involve *γ*-synchronization (e.g. a similarity in orientedness or texture in the absence of structural predictability, Figure 7). Further, global Gestalt principles do not necessarily involve *γ* (82; 83; 84; 85; 81). We also observed a parallel between dimensionality at the neuronal and image level, in that low-dimensional stimuli are mapped onto low-dimensional neural states (Figure 4). More specifically, when the image can be reconstructed by fewer spectral components (low dimensionality), neural activity is also dominated by energy in a relatively narrow frequency band (*γ*) as reflected by population-wide synchronized firing.

Oscillatory synchronization might reflect or modulate the way in which V1 neurons compute and transmit information, e.g. via resonance (86; 87). The E/I circuit generating V1 *γ* oscillations behaves like a linear resonator that filters stochastic input drive (88; 24; 89). Thus, for high stimulus predictability, spectrally broadband FF inputs into V1 may have a reduced impact. Conversely, the impact of lateral recurrent (V1 to V1) or even FB (V2 to V1) inputs that are also *γ*-rhythmic and carry predictive signals remains preserved (37). If *γ* affects the way in which different input signals are integrated, then that may well affect the variability and information content of V1 firing rate representations (37). Indeed, a close link between local V1 information processing and *γ*-oscillations is suggested by several findings; membrane potential fluctuations in the *γ*-frequency range are more strongly orientation tuned than the mean membrane potential (90), orientation tuning of firing rates in V1 fluctuates *γ*-rhythmically (91), and orientation tuning is strongly correlated with *γ* phase-locking (43; 91). Synchronization likely also has important consequences for the induction of synaptic plasticity (92; 28), which is mediated by the well-established mechanism of spike-timing-dependent-plasticity (93; 94; 28; 95; 92). Importantly, synaptic plasticity does not require *γ* to be present for each image and in a continuous manner, hence is compatible with the stochastic nature of gamma and the observation that it is only induced by a subset of image patches (Figure 4, (77)). Short-lived bursts of gamma oscillations could strengthen synaptic connections between neurons that, on average, predict each other’s visual inputs and thereby contribute to self-supervised learning of spatiotemporal natural-image statistics.

## Acknowledgements and Authorship Contributions

Conceptualization and design: CU, AP, MV. Data analysis: CU, MV. Design of neural networks and related methods: CU, MV. Implementation of neural networks: CU. Recordings: AP, CU, RR, JKL. Software: CU, AP, AL, KS. Surgery: JKL, WB, WS. Materials and reagents: WS, PF, MV. Supervision: MV. Writing: CU, AP, MV, with comments from the other authors. We thank Michael Schmid and Richard Saunders for implantation of monkey H. This projected was supported by ERC Starting Grant to MV (SPATEMP) and a BMF Grant to MV (Bundesministerium fuer Bildung und Forschung, Computational Life Sciences, project BINDA, 031L0167). The Titan X Pascal used for this research was donated by the NVIDIA Corporation.

## Methods

All procedures complied with the German and European regulations for the protection of animals and were approved by the regional authority (Regierungspräsidium Darmstadt).

### Surgical procedures

Three adult macaque monkeys (Macaca mulatta, two male, one female) were used in this study. Implantations were made in the left hemisphere of V1. In Monkey H, we implanted a Utah array with 64 microelectrodes (inter-electrode distance 400 *μ*m, half of them with a length of 1 mm and half with a length of 0.6 mm, Blackrock Microsystems), and inserted a reference wire under the dura towards parietal cortex. In monkeys A and I, we implanted a semi-chronic microelectrode array Microdrive (SC32-1, Gray Matter Research), containing 32 independently movable tungsten electrodes (inter-electrode distance of 1.5 mm); here, the microdrive chamber was used as the reference. Note that no histological verification of layers/depths was performed, because the animals are still alive. We estimate that our recordings mainly sample activity from layers 2-4, because the vast majority of recording sites do not show the typical inversion of the first deflection of the event-related potential as is found in the deep layers (96; 97), this is further supported by the electrode lengths in monkey H. Further details on surgical procedures can be found in (45).

### Behavioral task

All monkeys were trained on a fixation task. The animals were positioned 83 (monkey H) or 64 cm (monkeys A, I) in front of a 22 inch 120 Hz LCD monitor (Samsung 2233RZ, Ghodrati et al. (98); Wang (99)). Monkeys self-initiated trials by fixating on a small fixation spot, which was presented at the center of the screen, and had to maintain fixation during the entire trial. Trials during which the eye position deviated from the fixation spot by more than 0.8 (monkey H) or 1.44 visual deg (monkey I and A) radius were aborted. Correct trials were rewarded with diluted fruit juice. For further details on the task see Peter et al. (45).

### Recordings

Recordings were made with a Tucker Davis Technologies (TDT) system. Data were filtered between 0.35 and 7500 Hz (3 dB filter cutoffs) and digitized at 24.4 kHz (TDT PZ2 preamplifier). Stimulus onsets were recorded with a custom-made photodiode. Eye movements and pupil size were recorded at 1000 Hz using an Eyelink 1000 system with infrared illumination. For further details see Peter et al. (45).

### Visual stimulation

#### Image selection

Natural images were acquired from the Yahoo Flickr Creative Commons 100 Million (YFCC100M) Dataset (100). The resolution of these images was high enough to match the resolution of the LCD monitor given the stimulus size (see below). Images were included if user tags included any of the following: Animal, building, closeup, flower, house, indoor, landscape, natural, object, texture, tool, toy, tree. Any of the following tags led to exclusion: Blur, blurry, bokeh, digital, art, artwork, artist, text, writing, drawing, painting, cartoon, graphic, graphic+design, illustration, logo, desktop. A set of 340 images was selected as stimulus set for the monkeys. Note that from the same Yahoo Flickr database, a training and validation set was selected for the deep neural network training (see further below); this training set was different from the images presented to the monkey. All images were converted to grayscale, except for a subset of sessions in which we presented the images also in color. The 340 selected images had to fulfill the following criteria, which are comparable to the ones used in Coen-Cagli et al. (13): (1) A mean RGB value between 40 and 200. (2) An average luminance-contrast, measured as rootmean-squared (RMS) contrast, above 0.2. (3) A centroid of spatial frequency (defined further below in the Image Statistics section) greater than 0.5 dva (degrees of visual angle). These criteria excluded images that were excessively bright, dark, or spatially uniform. However, note that we studied V1 responses to uniform grayscale surfaces in (45)).

#### Image standardization

We cropped each image to 600×600 pixels, and applied two further transformations to the images, similar to (13): (1) We set the global luminance-contrast for each image to 0.6, by using a sigmoid projection of pixel values. (2) We rescaled and centered the images to have a mean RGB value of 128, equal to the background. For stimulus presentation, we approximately centered the images on the cluster of recorded V1 RFs. Stimuli had a width of 11.5 dva (monkey H, stimulus centered horizontally at x=+2 and vertically at y=-3 dva from fixation) or 15 dva (monkey A x=+3.55 and y=-0.12, monkey I x=+3.45 and y=-0.02).

#### Further image selection criteria

Stimulus sets for a given recording session consisted of 20 images, each presented for a total of 10-20 trials. Each session contained a subset of ten images where at least part of the image had a high spatial predictability (>0.85, see below). This was done to ensure that a sufficient number of image patches with high predictability were sampled, given the low probability of finding image patches with very high spatial predictability (Figure Extended Data 10). These images were found as follows: (1) Around the clusters of RFs, a region of 3 x 3 degrees visual angle was selected (ROI). (2) The ROI was divided into 1 degree patches and the spatial predictability of 9 non-overlapping patch locations were quantified (see section Image Statistics). (3) At least 2 out of 9 patch locations were required to have a predictability value (defined with SSIM) above 0.85. The other 10 images were, in terms of predictability, selected randomly. Note that correlations between spatial predictability and synchronization were also found below 0.85 (Figure 3), and that for all 20 images, there were typically some recordings sites with RFs corresponding to low predictability. Additionally, all 20 images were required to have, inside the ROI, an average luminance-contrast above 0.15 and a centroid of the spatial frequency below 8 cycles per degree; this prevents aliasing, given the visual acuity of macaques at the recorded eccentricities.

### Data analysis

#### Preprocessing

We analyzed only correctly performed trials. We downsampled the LFPs to ≈ 1.02kHz using Matlab’s decimate.m function. Line noise was removed using a two-pass 4^th^ order Butterworth bandstop filters between 49.9-50.1, 99.7-100.3 and 149.5-150.5 Hz. Similar to previous studies (101; 102; 103; 104; 45), we computed MU (Multi Unit) signals from the broadband signal, by bandpass filtering between 300 and 6000 Hz (4^th^ order butterworth), rectification, and applying low-pass filtering and downsampling in the same way as for the LFPs. For the calculation of rate modulations, this MU signal was smoothed with a Gaussian kernel with an SD of 20 ms. For further details see Peter et al. (45).

#### RF estimation

RF were mapped with moving bar stimuli (spanning the entire monitor). Moving bars (width 0.1 deg, speed 10-17 deg/s) were presented with 8-16 different orientations. MU responses were computed as a function of stimulus position, after correcting for response latency. MU responses for each movement direction were then fitted by a Gaussian function. We used this fit to extract the 10th percentile and the 90th percentile. Across the 16 directions, this yielded 32 data points, which were fit with an ellipse. This ellipse was defined as that MU’s RF. The RF size was defined as the diameter based on 2*sqrt(a*b), where a and b are the major and minor radius. The preferred stimulus orientation was also computed using these bar stimuli (and was highly consistent with orientation tuning based on static gratings).

#### Electrode selection

We included all electrodes for analysis that met the following criteria: (1) The MU showed a response to RF stimulation that was at least two SD above stimulation outside the RF. (2) The MU response during the response period (0.05-0.15 s post stimulus onset) of at least one stimulus condition in the session was at least 2 SD above the corresponding baseline (−0.4-0 s before stimulus onset).

#### Estimation of LFP power spectra

For monkey H, we made use of the constant spacing of neighboring electrodes in the array to improve power spectral estimation: Power spectra were approximated by the complex mean of the cross-spectra of a channel with its two same-depth nearest neighbors. This reduced uncorrelated high-frequency fluctuations due to spiking, which can affect the 1/ *f* slope of LFP spectra; qualitatively similar results were obtained using the unipolar power spectra. LFP epochs were multi-tapered (± 5 Hz smoothing) (42), Fourier transformed and squared to estimate LFP power spectral densities.

#### Quantification of LFP gamma-band and beta-band amplitude

In Peter et al. (45), we developed an algorithm to extract the amplitude of narrow-band gamma-band (or beta-band) oscillations, with several advantages compared to previous methods. Note that qualitatively similar correlations with spatial predictability were obtained by simply using baseline-corrected power in *β* and *γ* bands (see Figure Extended Data 3, however baseline-corrected power can be skewed by firing rates which results in a stronger correlation with luminance-contrast (Figure Extended Data 20). The algorithm had the following structure:

1. Power spectra were log-transformed and the frequency axis was also sampled in log-spaced units to avoid over-weighing the contribution of high-frequency datapoints. All subsequent polynomial fits were performed on the 5-200 Hz range.
2. 1/F^n^ correction was performed by fitting an exponential to the LFP power spectrum, excluding data points in the range of 10-85 Hz in order to avoid any influence on peak detection. (For a subset of figures, namely 3 and 5, we corrected the power by dividing by the pre-stimulus baseline).
3. To determine the polynomial order, we used a cross validation procedure to prevent overfitting. Polynomials of order 1-20 were fit to Δ*P* as a function of frequency for a “training set”. We then evaluated the mean squared error using the same polynomial fit on a “test set” for each of the 20 orders. This procedure was then repeated for multiple iterations and we chose the order with the best median performance.
4. On the polynomial fit, local maxima and minima in the *β* (18-30 Hz) or *γ* (30-80 Hz) range were identified. The peak frequency was the location of the maximum. The amplitude was then assessed from the difference between the value of the polynomial fit at the maximum and the average of the polynomial fit at *F*_*min*_ and 2*F*_*max*_ − *F*_*min*_, where *F*_*min*_ is the frequency of the nearest local minimum to the left of the maximum (we used the left one, to avoid any influence of spike-bleed-in at higher frequencies).
5. The amplitude was quantified as a fold-change.

#### Rate modulation

Rate modulation was computed as

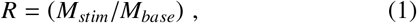

where *M*_*stim*_ and *M*_*base*_ represent the MU firing activity in the stimulus and baseline period, respectively. Spike-density functions were normalized in the same way.

#### Statistics

Error bars or shaded error regions correspond to ± one standard errors of the mean (SEM) across recording sites. Violin plots show the median together with the 25-75 percentiles and the data distribution estimated using Matlab’s ksdensity.m function. The statistics used in the rest of the manuscript were as follows:

- In Figure 3C, 5B, and 6A, we show average neural activity (72 recording sites) as a function of spatial predictability or luminance-contrast. We pooled all 72 channels and RF image patches together and then formed non-overlapping bins of 250 RF image patches. We then performed a Spearman-rank correlation across these 250 images.
- In Figure 3D and 5C, we correlated *γ*-peak amplitude with predictability or luminance-contrast across sessions for each channel separately. We tested whether the average correlation was significantly different from zero across 72 channels by using a two-sided T-test.
- In Figure 5C and 6D, we tested whether different models were significantly different from each other by using 10-fold cross-validation, and a paired T-test across 72 recording sites. The additional regression variables used in 5C were: Center luminance, center spatial frequency and absolute deviation from preferred MU orientation, as well as the luminance and spatial frequency of the 224×224 pixel image patch surrounding the RF image patch.
- In Figure 3E and Figure Extended Data 11, we performed a two-sided T-test between nature and man-made or object-boundary vs. no boundary categories (N=72 recording sites).
- In Figure 6E, we used a Pearson correlation to quantify the trend across layers.
- In Figure 1I, we compared the strength in correlations between blocks of VGG-16 layers using a pairwise two-sided T-test across N=72 recording sites (early: layer 1-4; middle: layer 5-9; deep: 10-13).
- We repeated 5 experimental sessions twice with 5 different stimulus sets with 100 stimuli. Using this we estimated the variance of the estimated means (which mainly results due to limited number of trials) and corrected for this variance.

### Identification of object boundaries and man-made vs. nature pictures

To automatically identify object boundaries, we used the Mask-R convolutional neural network. Using this network, we determined if there is a detected object contour that intersects with the MU’s RFs. We distinguished man-made vs. nature picture as follows: Pictures were categorized as man-made if man-made structure (e.g. buildings) was within the 4 × 4 degree image region centered on the RFs of the recording array (for examples see Figure 3). Images with both nature and man-made content were not considered for this comparison.

### Deep Neural Networks methods

We used DNNs for several purposes. In Figures 2–6 we trained a network to predict missing parts of an image to obtain several measures of spatial predictability. We will first describe the stimuli used to train this network, its architecture and training procedures. Following this, we describe other networks in which we directly predicted neural activity from the image.

### Preprocessing of training stimuli

We resized the images from the Yahoo Flickr dataset (see Section “Visual Stimulation”) to 300×300 pixels. To improve robustness and generalization, we applied standard data augmentation of the images for the training, consisting of several operations from the Tensorflow image module (105). Each operation had a 50% chance of being applied for a given image: brightness (max_delta=0.1), contrast ([low, high] = [0, 1]), hue (max delta = 0.1), saturation ([low, high] = [0, 1]), convert to black & white, horizontal flip. Resulting images were then randomly cropped to 224×224 pixels. The mean RGB value [123.68, 116.779, 103.939] of ImageNet was subtracted from each image, in order to use the VGG-16 network for the initial layers of the network (106).

#### Mask generation

In order to make a network that can robustly predict missing inputs, we trained the network with binary masks that either had occluders or missing pixels (low signal-to-noise ratio). Binary masks were randomly selected from 3 types: elliptic, rectangular, or salt and pepper noise. Rectangular and elliptic mask types consisted of 2-3 missing regions that were randomly selected to be 20×20 pixels to 80×80 pixels (0.5-2 degrees) in area. One side of the rectangle (or axis for the ellipse) was randomly picked between 20 to 80 pixels. Finally random rotations were applied to each missing region. Salt & Pepper noise had a sparsity ratio of at least 20%. Images including the mask were randomly cropped to size 224×224, and horizontally flipped with a 50% chance.

### Architecture of deep neural network for inpainting

For the inpainting we relied on Deep Neural Networks. Note that qualitatively similar correlations (however of smaller magnitude) were obtained when generating image predictions based on an algorithm that does not use deep neural networks (107), demonstrating the generality of our approach (Figure Extended Data 21). The neural network architecture was based on the U-Net architecture (108; 109), with the following modifications: For initialization, the encoder part of U-Net was replaced by all the convolutional and pooling layers of the VGG-16 network, using the Keras implementation (106; 110). Transfer learning using VGG-16 has been previously used in image segmentation (111), image reconstruction (112), style transfer (61), and image inpainting (113). The resulting network architecture consisted of five blocks, each of which had two or three convolutional layers (3 × 3) with ReLU (rectified linear) activation functions, followed by a max-pooling (2×2) operation. The decoder consisted of five blocks, each with a nearest-neighbor upsampling layer (2×2), followed by two convolutional layers. The output layer was a convolutional layer with, as is conventional, a linear activation function.

#### Partial Convolution

All convolution operations in the network, including the VGG-16 network, were implemented as partial convolutions. Partial convolution has been introduced with the sparsity-invariant convolutional network where the input to each convolution is paired with a binary mask indicating which pixels are observable or missing, respectively (112). Partial observability of the inputs during the training makes the network robust to input sparsity, regardless of the task of the network. We implemented a modified version with mask updates per network operation, as described in (113). The idea of partial convolution is that the missing region is gradually filled, and that the filled-in information is used for filling in the rest of the missing pixels in an iterative way.

### Loss function for training

Convolutional neural network (CNN) activations have been previously used as a basis for perceptual similarity metrics instead of traditional measures such as SSIM (114) or L2-norm (115; 61; 60). Even though these perceptual loss functions based on CNNs match human perception better, image generation based on those loss functions can suffer from high frequency artifacts (116; 117). To minimize high-frequency artifacts, we therefore implemented the reconstruction and content loss functions in the Fourier domain, similar to decorrelated image parameterization described in (118; 119).

Total loss consisted of reconstruction, content, and style losses:

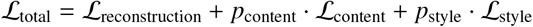

Here, *p*_style_ and *p*_content_ are hyperparameters (see further below). Reconstruction loss consisted of three terms: (i) The difference between the amplitudes of Fourier transforms; (ii) the log difference between the Fourier amplitudes; (iii) the phase similarity between the predicted and the original image, which has been previously applied to auditory signal synthesis based on the Fourier space (120):

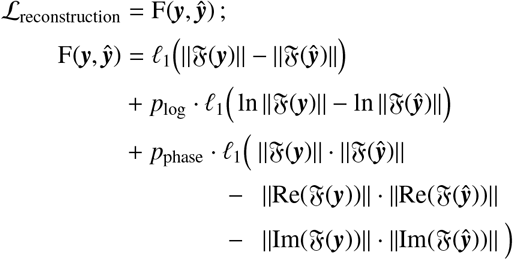

Here, ***y*** and ***ŷ*** are the original and predicted image. The operator 𝔉 denotes the Discrete Fourier Transform, ‖𝔉 (***y***) ‖ denotes the Fourier magnitude, and 𝓁_1_ denotes the L1-norm across all frequencies.

Content loss was also defined in the Fourier domain, taking as input the *c*th AN activation in the *λ*th layer activations of the VGG-16 network:

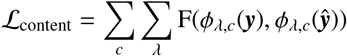

Finally, style loss is the difference between the Gramian matrices of the Fourier amplitude of the predicted and the original images, where the Gramian matrices contain the covariance matrix across AN activations:

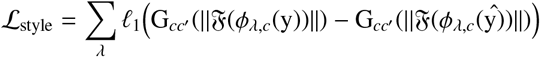

Four layers of VGG-16 network with fixed weights were used for content and style loss.

### Training and Hyperparameter Optimization

We initialized the weights of the encoder as VGG-16 model weights that were pre-trained on ImageNet. The remaining weights were initialized using He-initialization (121) and bias terms as 0. The network weights were optimized using the Adam optimizer with a learning rate of 5*e*^−5^, *β*_1_ = 0.9, *β*_2_ = 0.999, *ϵ* = 1*e*^−7^, (122). All hyperparameters were defined as continuous variables where the search space was [1e-5, 1e3] sampled in logarithmic scale (123). We optimized the hyperparameters using a method that combines Bayesian optimization with Hyperband as described in (124). We ran the optimization for 50 iterations with a minimum budget of 8 and a max budget of 16. For hyperparameter optimization, we used a subset of the image dataset for training (16384 images) and validation (1024 images) of the network. We used SSIM based reconstruction loss as the evaluation metric for the hyperparameter optimization. To analyze the hyperparameter space and the importance of individual hyperparameters, we usd fANOVA (125). The importance of a hyperparameter is the fraction of explained variance (mean across 100 repetitions ± SEM) of the validation SSIM-loss across the entire hyperparameter space. The resulting hyperparameters and their importances were:

| Name | Value | Importance |
| --- | --- | --- |
| $p_{style}$ | 0.18 | $0.13 \pm 0.002$ |
| $p_{content}$ | 36.90 | $0.22 \pm 0.002$ |
| $p_{log}$ | 5.54 | $0.14 \pm 0.002$ |
| $p_{phase}$ | $3.02e-05$ | $0.29 \pm 0.003$ |
| $p_{lr}$ | $4.48e-05$ | $0.14 \pm 0.002$ |

### Image Statistics

#### Spatial Predictability

for a given image, predictability was computed by masking out the central 1 degree patch (similar in size to previous studies studying contextual modulation, Coen-Cagli et al. (13)). The RF image-patch was then predicted by the inpainting algorithm described above. The to-be-predicted image patch is denoted **I**, a matrix of *N* × *N* pixels.

#### Structural predictability

Structural predictability was defined as the squared Pearson correlation of two images or average correlation across VGG-16 AN activations. This was defined for each layer *λ* of the VGG-16 separately:

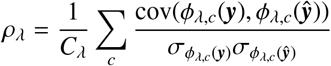

#### Content predictability

Content loss was defined as as the L2-norm of the difference between the VGG-16 *λ* layer AN activations of two images. This was defined for each layer of the VGG-16 separately:

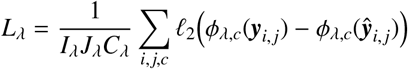

#### Luminance-contrast

Luminance-contrast was measured as the Root-mean-square (RMS) contrast:

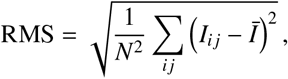

where *Ī* is the mean value of *I*. RMS was defined to range between 0 (minimum) and 1 (maximum) luminance-contrast.

#### Spectral Centroid

The center of mass of the power spectrum is the mean spatial frequency in the image patch, *f*_*k*_, weighted by the total power, *r*_*k*_ in the frequency bands.

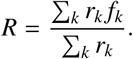

#### Orientation

Mean orientation in the RF was computed weighted by the total power in the orientation band [0:5:180] degrees, across spatial frequencies 1.0, 2.8284, 8.0 cycles per degree, similar to the orientation selectivity of the receptive field. We estimated the orientation selectivity of the RF as the weighted circular mean across different orientations as described in (126).

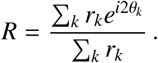

*R* is the population resultant vector, *r*_*k*_ is the peak MU response for the orientation *θ*_*k*_. Orientation selectivity magnitude is given by |*R*| and the mean angle is given by arg(*R*).

#### Compressibility

Compressibility was defined as the negative bits per pixel (bpp) which is commonly used to benchmark image compression methods. For image compression, we used a context adaptive entropy based deep neural network model that outperforms the traditional image codecs such as BPG or JPEG, as well as other previous DNN based approaches (127). For each RF image patch and its surround (224×224 pixels), we compressed the image using the MS-SSIM optimized model with the quality level set to 5 to get a bits-per-pixel (bpp) measure of the compression. For the same image patches we computed the average predictability across 52×52 pixel non-overlapping sub-regions of the whole patch.

#### Dimensionality

Dimensionality was determined by first taking the two-dimensional fast Fourier transform of the RF image-patch, taking the rotational average, and ranking the spectral components by magnitude. Dimensionality was then defined as the slope of the resulting spectrum.

#### Homogeneity

We computed P(homogeneity) using the inference model described in (128) similar to (129; 13). This model consisted of 72 filters with 4 orientations (0, 45, 90, 135 degrees) and 2 phases (even, odd-symmetric quadrature pairs) each, at 9 locations (center and 8 surround locations circular around the center RF with a radius of 6 pixels). Center and surround RFs had a diameter of 9 pixels and peak spatial frequency of 1/6 cycles/pixel. We trained the model with our natural image dataset, downsampled by a factor of 0.22 to match the model RF size (9 pixels) using the code provided by the authors(130). P(homogeneity) was computed as 1-P(heterogeneity), where P(heterogeneity) is the average inferred probability of hetereogeneity of 4 center units which was the output of the inference model.

### Human perceptual similarity

We used the Berkeley Adobe Perceptual Patch Similarity Just Noticeable Distance (JND) dataset to evaluate content and structure losses as human perceptual similarity metrics. In the JND experiments, participants were presented two image patches for 1 second each, with a 250 millisecond gap in between. They were asked if the patches were the same or not. In order to evaluate content and structure perceptual similarity metrics, we used the layer specific VGG-16 AN activations of the original and distorted image as predictors for a logistic regression classifier to predict human perceptual similarity judgments. We calculated the Area Under Curve (AUC) to quantify how well the different VGG-16 layers and similarity metrics (LPIPS, LPIPS structure, SSIM, structure) explain human perceptual similarity judgments.

### Predicting neural activity from VGG-16 activations

In Figure 1, we predicted neural activity from the VGG-16 AN activations. If VGG-16 ANs had a smaller RF than 1 dva, then only VGG-16 ANs were used with a RF center within the 1 degree region around the multi-unit’s RF center. If VGG-16 ANs had a larger RF than 1 dva, then only VGG-16 ANs were used that fully covered the central 1 degree. For each location in the VGG-16, we predicted the neural responses from a vector of VGG-16 ANs with different feature selectivities. For this, we used 10% of the stimuli as a test set and used the rest as the training set. We used L1-constrained linear regression and 10-fold cross validation to select the L1-constraint parameter *λ*. Regression coefficients that best explain the training set were then used to predict the neural signals for the test set. The correlation values (r) were averaged across VGG-16 locations. Receptive field sizes in the VGG-16 are shown in Table 1. We calculated the receptive fields for VGG-16 ANs as described in (131). The receptive field sizes of the neural network in the middle layers were comparable to the receptive field sizes in V1, which has been previously described in Cadena et al. (46).

### Tools

Neural network visualizations were made using PlotNeuralNet (132). Software and packages used for the analysis include MATLAB, matplotlib (133), numpy (134), docker (135), tensorflow v1.14 (105), and (110).

## Data availability

The data that support the findings of this study are available from the corresponding author M.V. upon reasonable request.

## Code availability

Data were preprocessed and analyzed using the FIELDTRIP Toolbox (136) and an algorithm, as described in the Methods and previously published in (45). Custom software developed for Figures 2–6 and 8 are made available on the github repository (https://uranc.github.io/U-Net-Pred/, including small demos. All other used software is referenced in the text when used and available online.

---

**Figure Extended Data 1:**
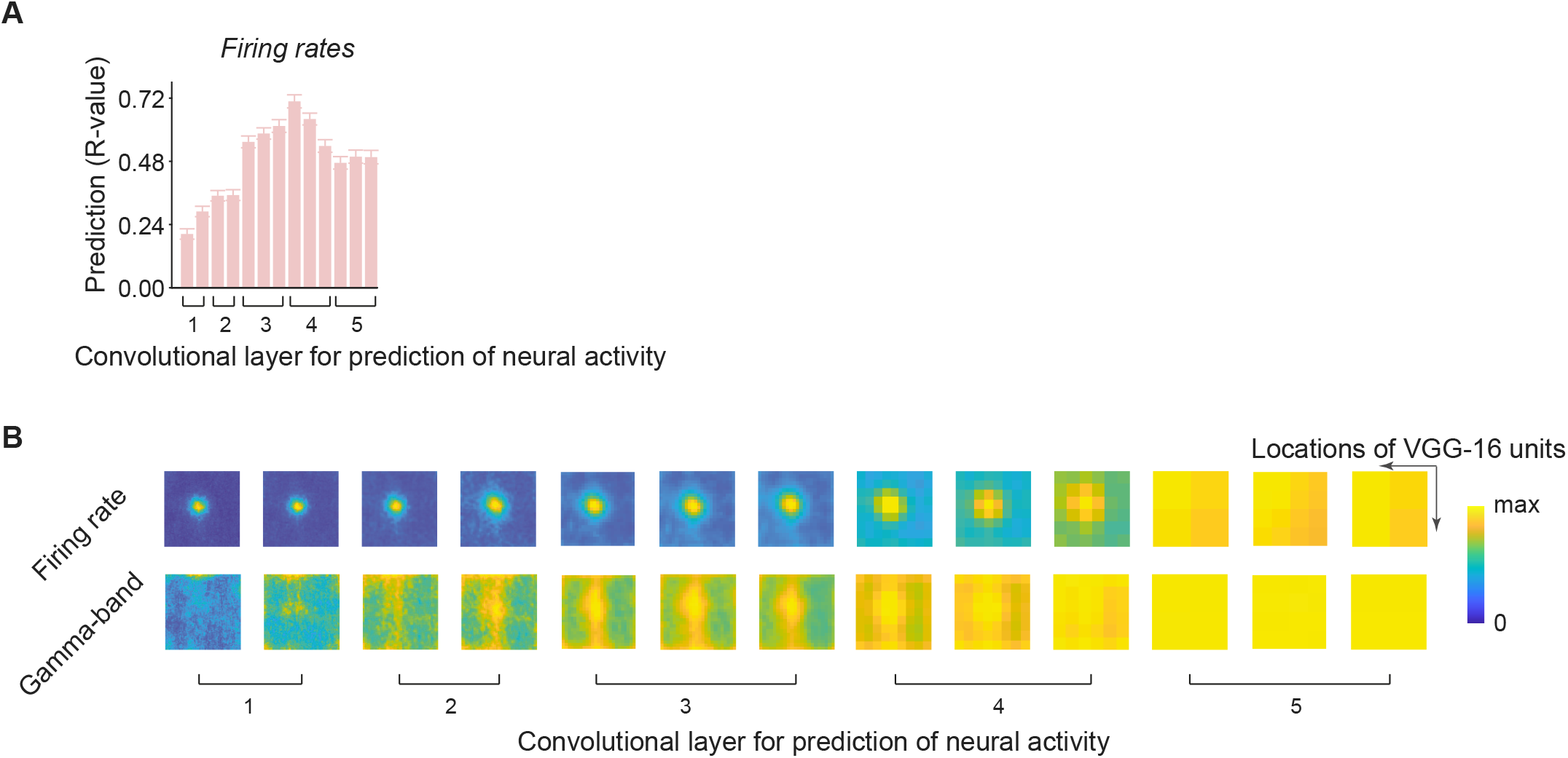
**(A)** Prediction of early firing rate intensity with a convolutional neural network for object recognition (VGG-16). Same as Figure 1I, but now for early rates. **(B)** Prediction accuracy depending on the RF location of VGG-16 ANs in the image. In this case, we predicted neural activity from all units in a 3×3 image using sparse L1-regression. Shown are the prediction weights, which reveal circular RFs for firing rates already in the earliest layer.

**Figure Extended Data 2:**
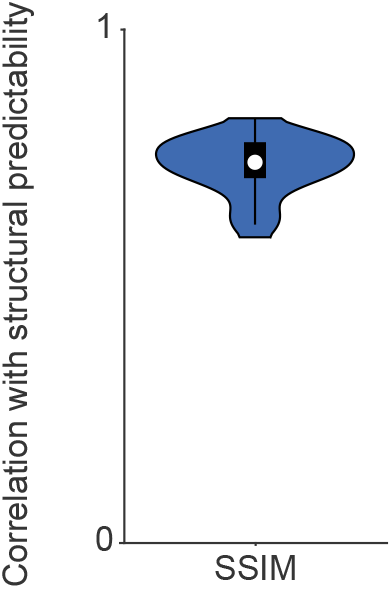
Spearman correlation of the perceptual similarity measure SSIM to the structural predictability measure defined in Figure 3 across images.

**Figure Extended Data 3:**
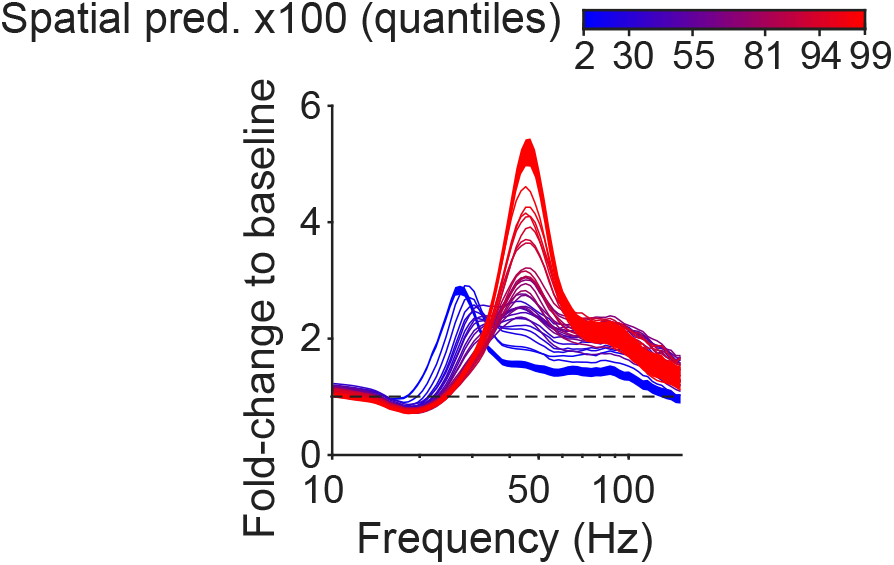
Average baseline-corrected LFP power spectra. SEMs are shown for the lowest and highest quantile of predictability.

**Figure Extended Data 4:**
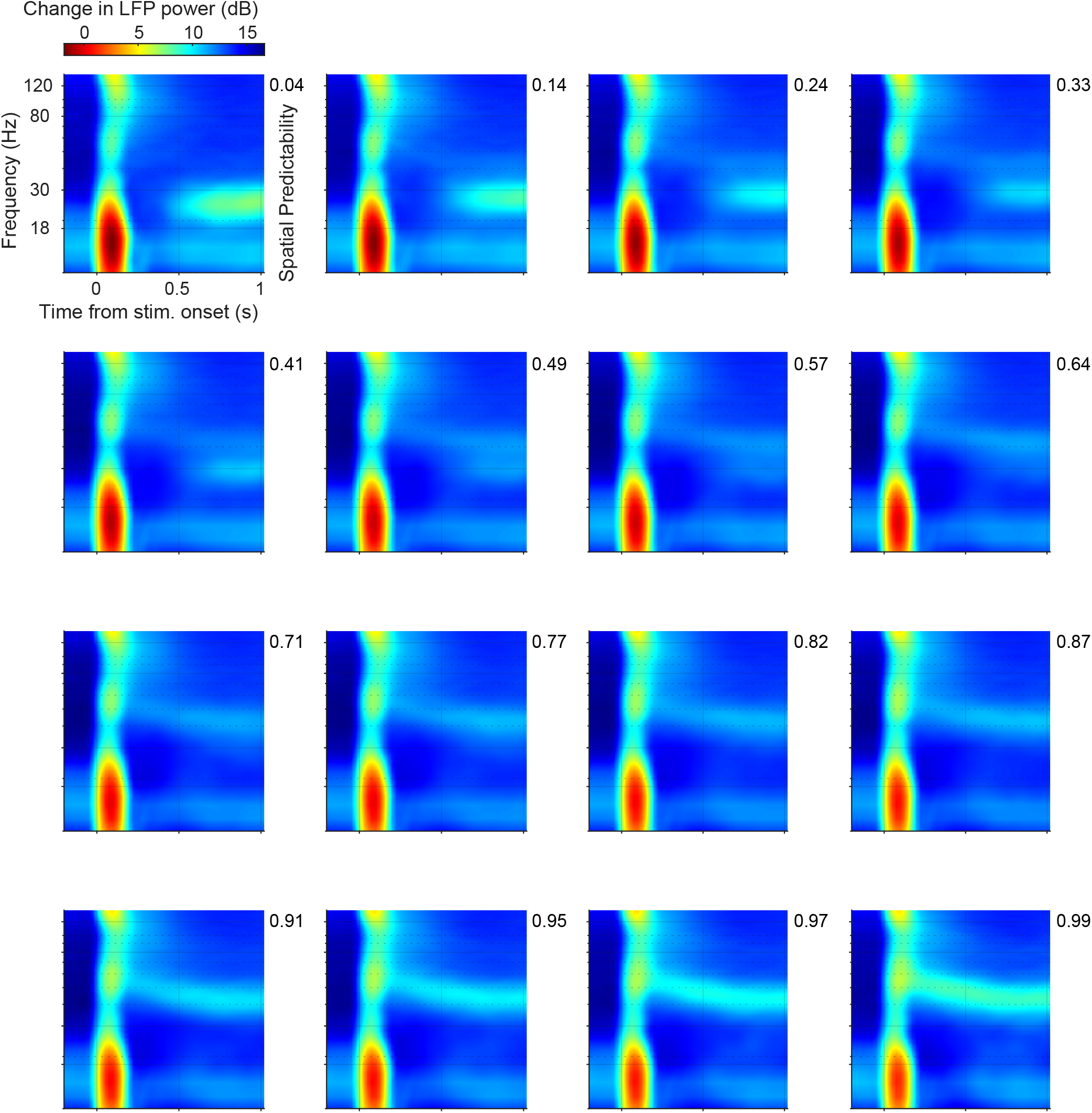
Time-frequency representations for 16 different levels of predictability (left-to-right, top-to-bottom). Time-frequency representations were computed using discrete prolate spheroidal sequences (multi-tapers for +/-5 Hz smoothing) with a 300ms time window sliding in steps of 50ms.

**Figure Extended Data 5:**
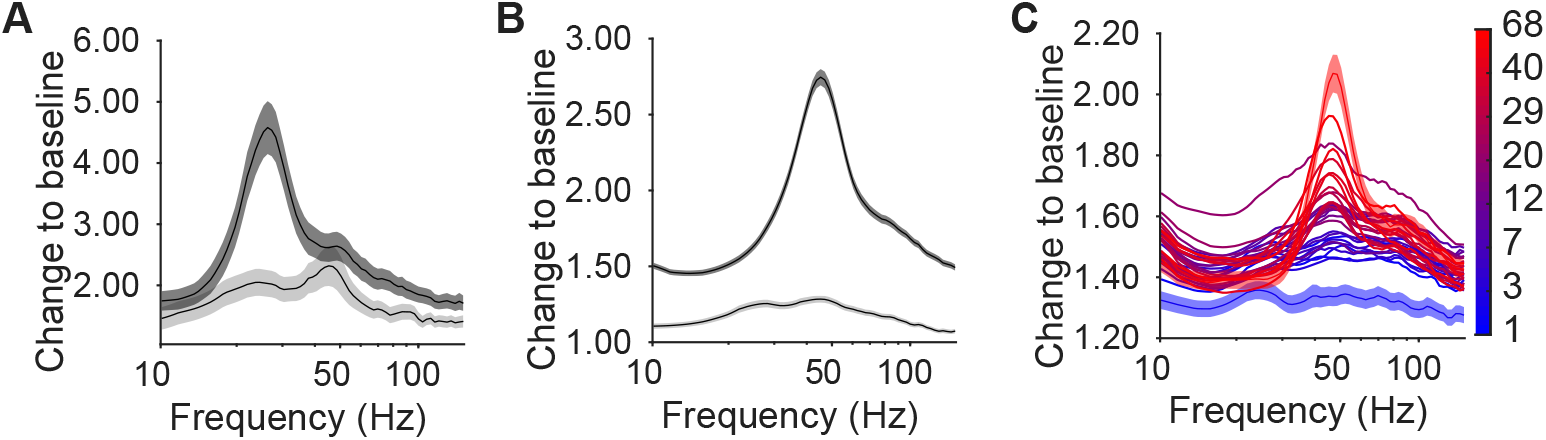
**(A)** Average MUA power spectrum for image combinations (across sessions) with 25% highest (black) and 25% lowest *β*-peaks (gray). Like the LFP, the MUA shows rhythmic fluctuations in the beta-band with a peak around 26 Hz. **(B)** Same as (A), but now for *γ*. **(C)** MUA power as a function of the product of structural predictability and luminance-contrast, as shown in Figure 5C. Blue-to-red colors indicate quantiles of contrast × predictability. MUA power shows a gradual increase in *γ* amplitude. All panels are for monkey H.

**Figure Extended Data 6:**
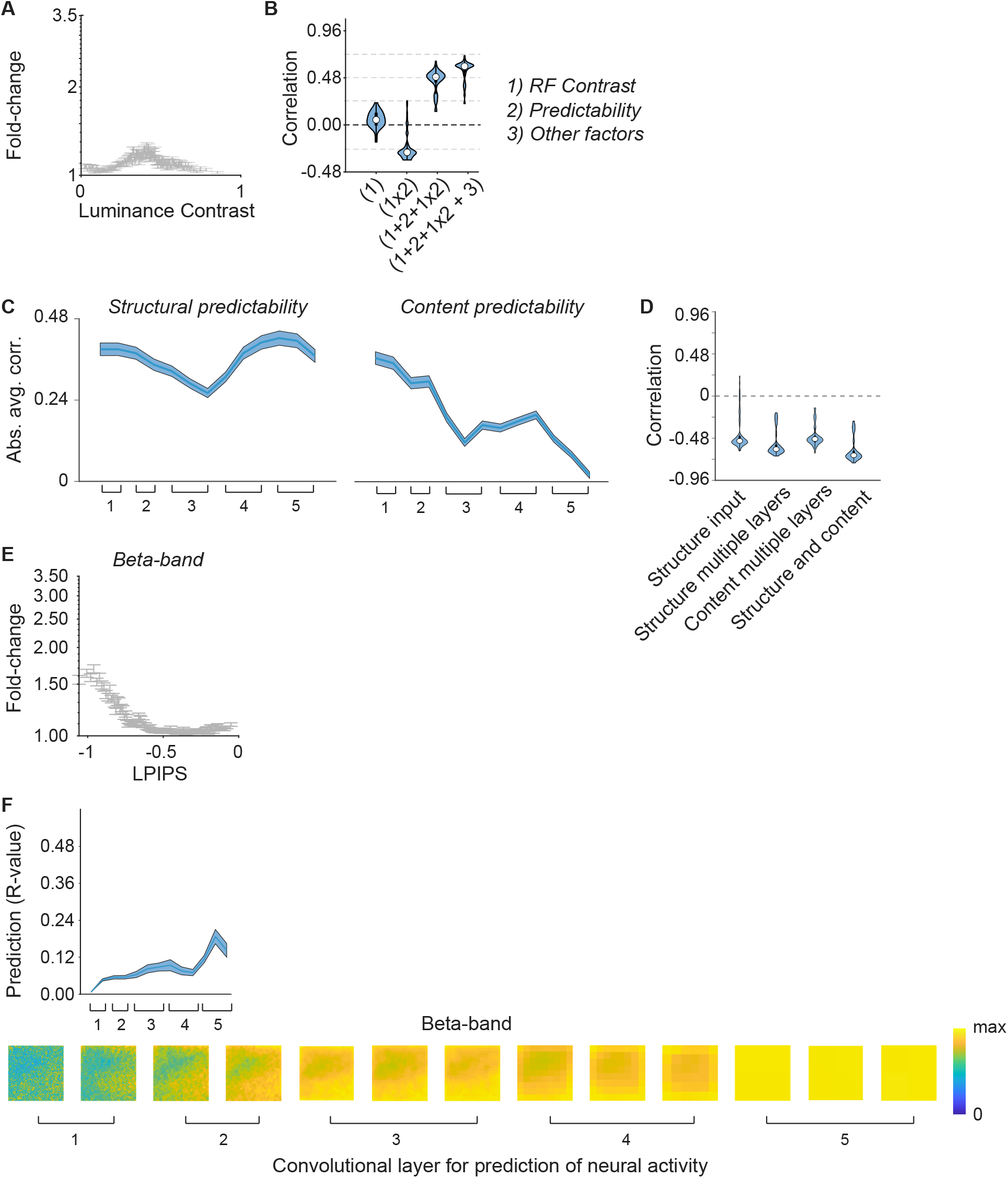
Main results for *β* synchronization. **(A)** Average 1/f corrected *β*-peak amplitude for different levels of luminance contrast, similar to Figure 5B. *β* power peaks at intermediate luminance contrasts. **(B)** Similar to Figure 5C. Beta was very weakly correlated with luminance contrast and strongest for intermediate contrasts. Including other image factors as predictors led to a small increase in explained variance for *β* synchronization. **(C)** Correlation of *β*-power with structural (left) and content (right) predictability across VGG-16 layers. *β* showed a significant decrease in absolute correlation with predictability across layers for content (*R* = 0.9, *P* < 0.001). This is consistent with the idea that *β* synchronization propagates from higher areas to early visual cortex (137; 48; 34; 138). **(D)** Left to right: (1) Correlation with structural predictability at the input image level; (2) regression model based on structural predictability across layers 1conv2, 2conv2, 3conv3, 4conv3, 5conv3; (3) regression based on content predictability in those layers; (4) regression based on both content and structure predictability in those 5 layers. **(E)** Correlation of LPIPS with *β* synchronization across bins of LPIPS predictability (correlation was significant at *P* < 0.001). **(F)** Top: For *β* synchronization, we found that explained variance was also low (peaking around 0.20) compared to firing rates. Prediction accuracy increased with VGG-16 layer, and *β* synchronization was best predicted from the activity of units in the deeper layers of the VGG-16 (5conv2) (similar to Figure 1) (deep vs. early and middle VGG-16 layers: *P* < 0.001, paired T-test). Bottom: RFs for *β* synchronization did not show a localized RF structure that was related to VGG-16 units.

**Figure Extended Data 7:**
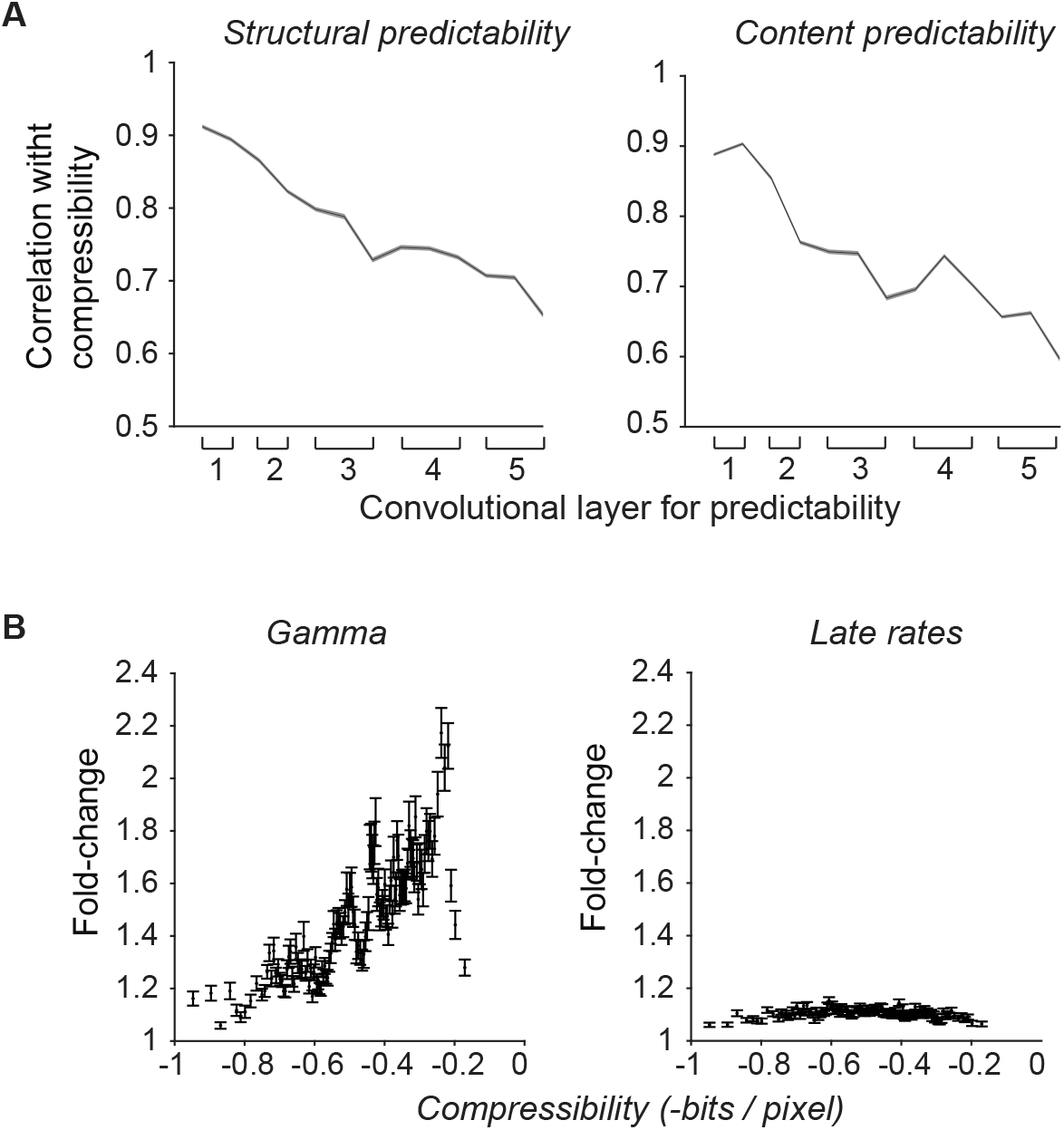
**(A)** Structural predictability and content predictability defined across different layers (Figure 6) vs. compressibility. Both correlations were negative (R=−0.97, R = −0.92, *P* < 0.001). **(B)** *γ*-synchronization and firing rates across bins of compressibility.

**Figure Extended Data 8:**
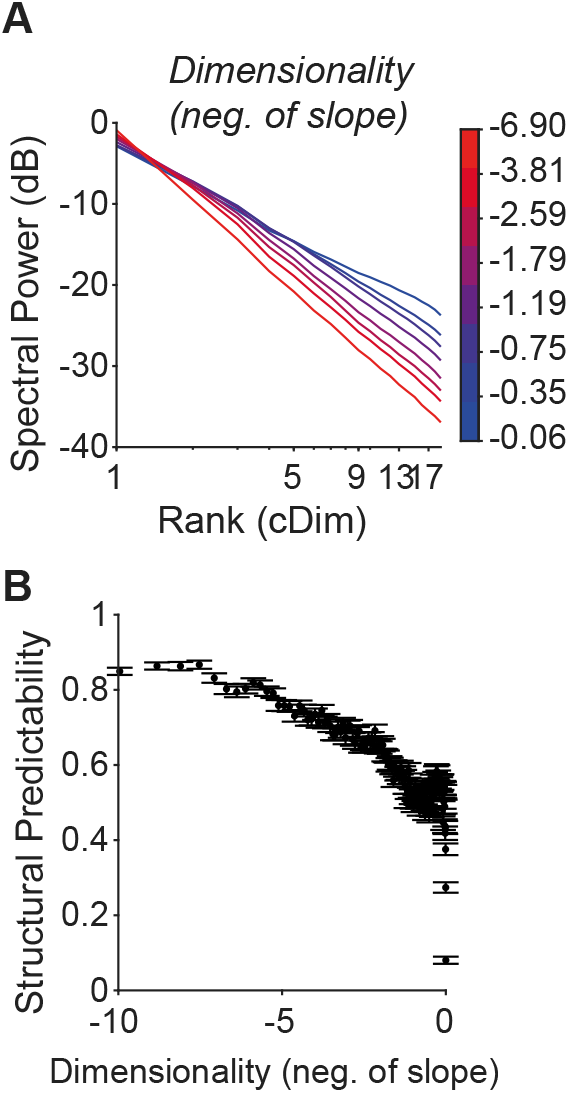
Dimensionality, predictability and *γ* synchronization. **(A)** Dimensionality was determined by first taking the two-dimensional fast Fourier transform of the RF image-patch, taking the rotational average, and ranking the spectral components by magnitude. Dimensionality was then defined as the slope of the resulting spectrum. An image with low dimensionality can be represented only by a few spatial frequency components. **(B)** Images with low dimensionality had high spatial predictability (R across bins = −0.91, *P* < 0.001).

**Figure Extended Data 9:**
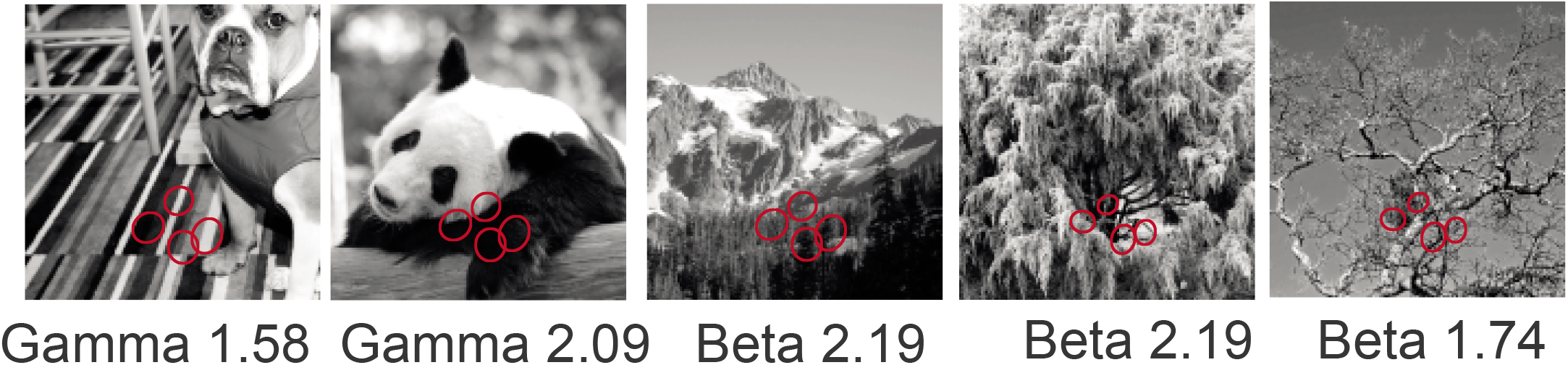
Example images with *β* and *γ* values for a set of RFs.

**Figure Extended Data 10:**
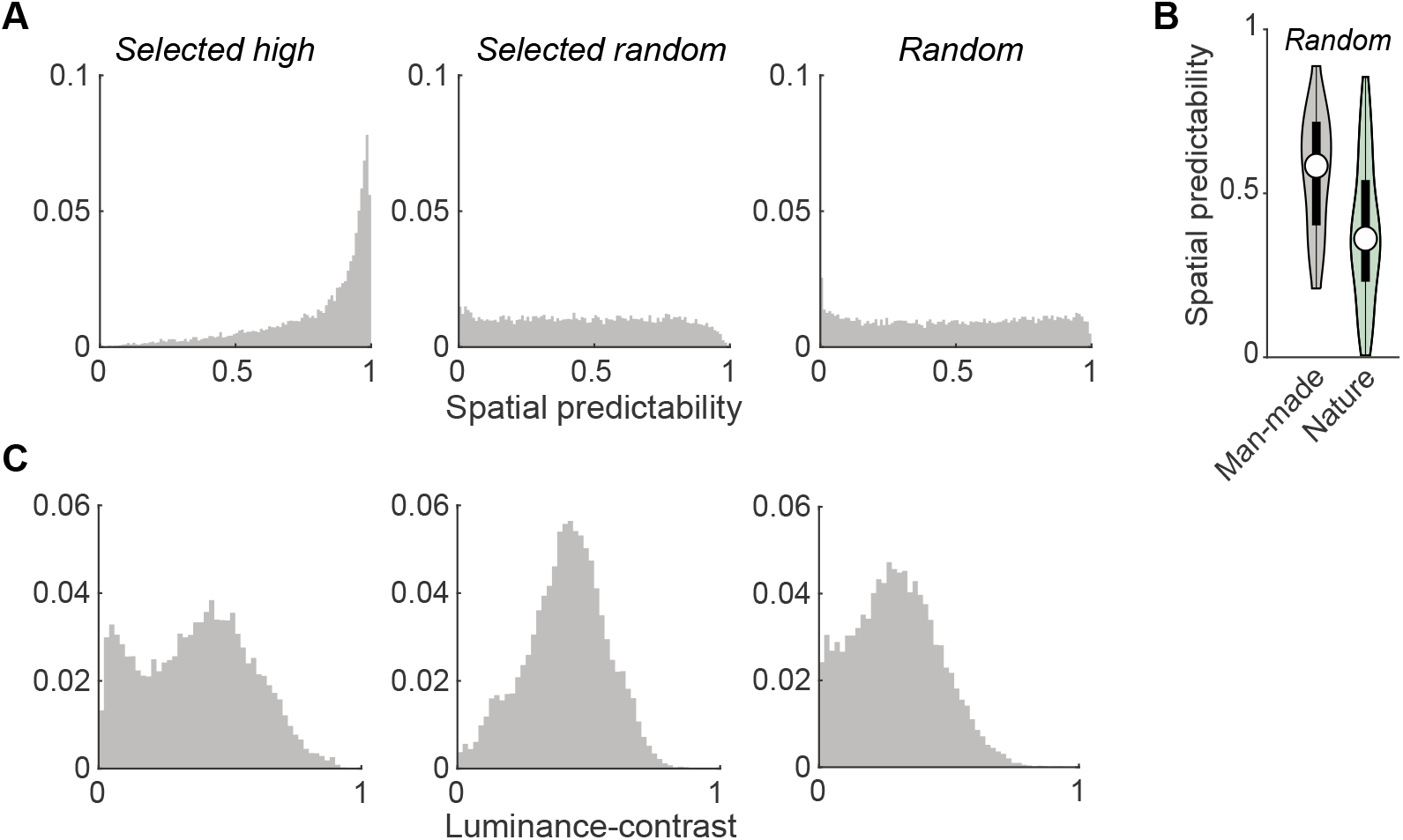
**(A)** Histograms of spatial (structural) predictability for three image categories. Selected high: 10/20 images that were randomly selected with additional criteria on predictability in the RFs of the neurons. Selected random: images that fulfilled the criteria as stated in the Methods section, but had no additional criteria on spatial predictability (10/20 images per session). Random: images that fulfilled the criteria on global luminance contrast (> 0.2), luminance (between 40 and 200) and spatial frequency (average centroid greater than 0.5 degrees of visual angle). **(B)** Average structural predictability for the random images category, separate for nature and man-made contents. Shown are the median, 25/75% percentiles and data distribution across images. Man-made images have higher spatial predictability (T-Test, *P* < 0.001). **(C)** Same as (A), but now for luminance contrast.

**Figure Extended Data 11:**
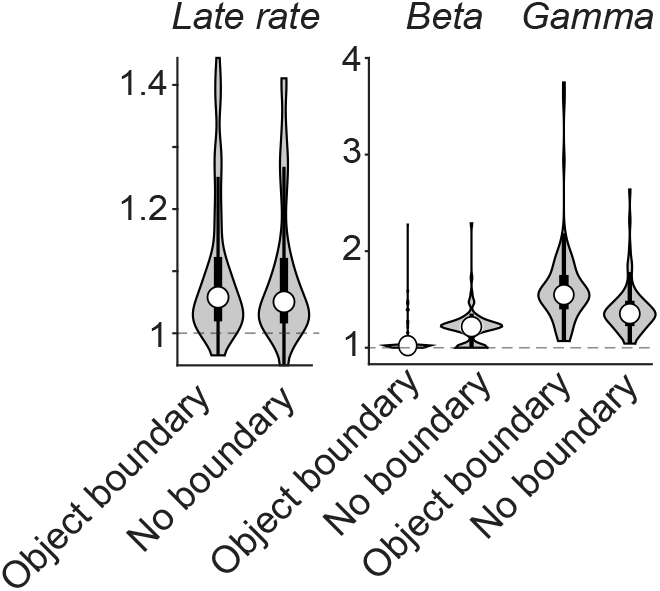
Fold-changes in neural activity for image patches that contain object boundaries versus image patches that do not. Comparisons were significant for *β* (*P* < 0.001) and *γ* (*P* < 0.001), but not for early and late rates (*P* = 0.94 and *P* = 0.73, T-test).

**Figure Extended Data 12:**
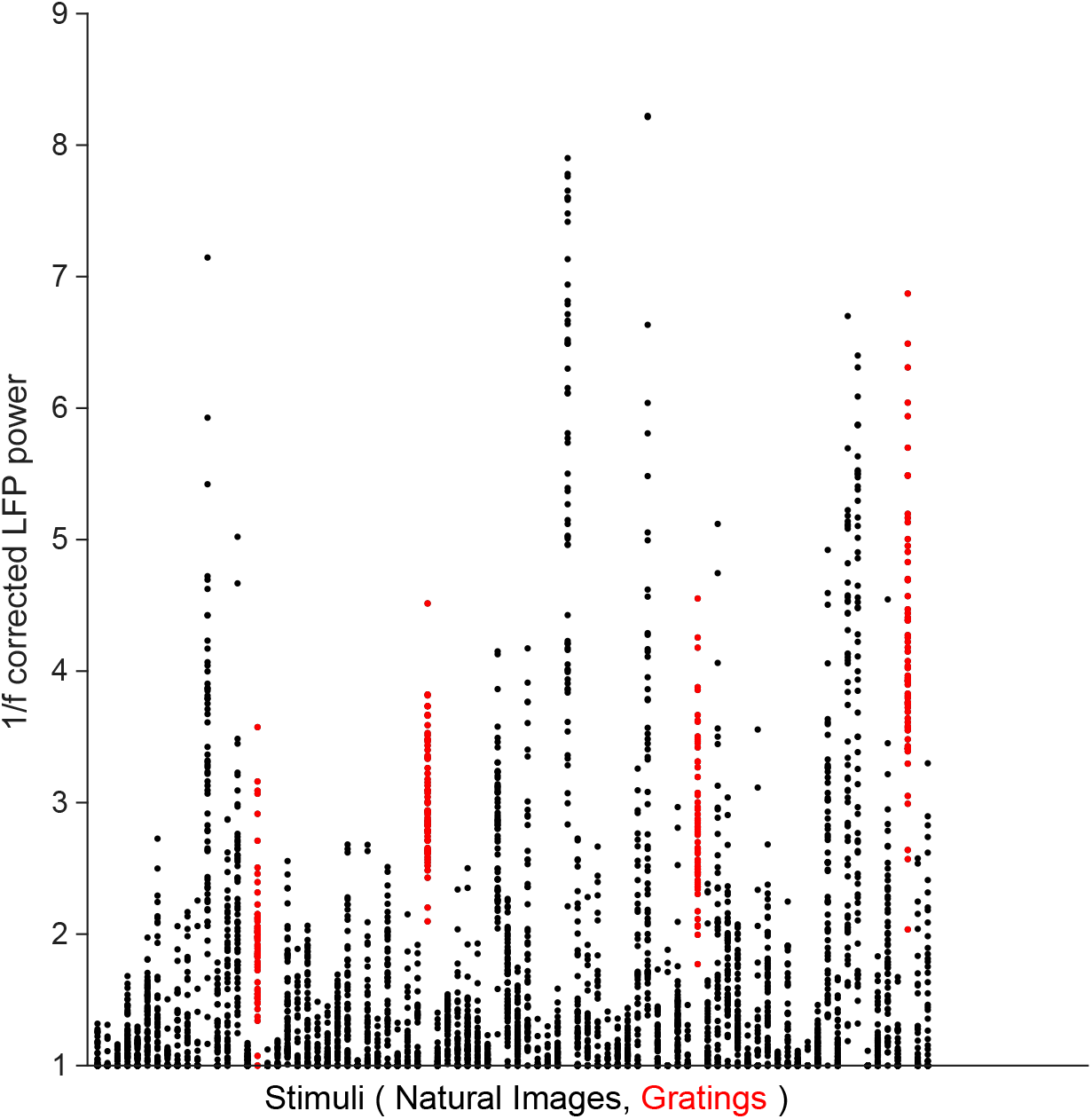
Gamma-band responses for 4 sessions that included a stationary grating stimulus at 2c/d spatial frequency, presented in vertical or horizontal orientation. Each dot represents a trial. Stationary gratings yield strong gamma, however several stimuli in those sessions induced stronger gamma-band activity.

**Figure Extended Data 13:**
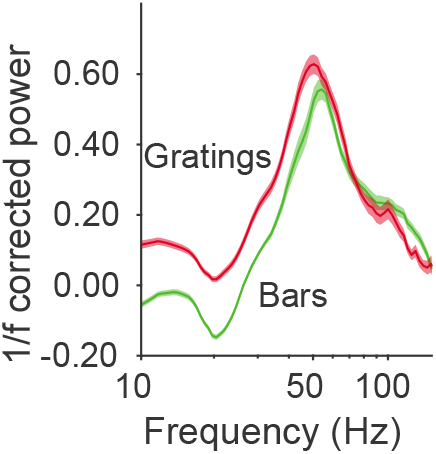
Bar stimuli generate comparable *γ* response to gratings. Square-wave grating stimuli presented were 6 degrees in diameter with a spatial frequency of 0.5,1,2,4 c/d. Bar stimuli were matched in width to the respective spatial frequency. Average across 28 channels from monkey H and monkey A.

**Figure Extended Data 14:**
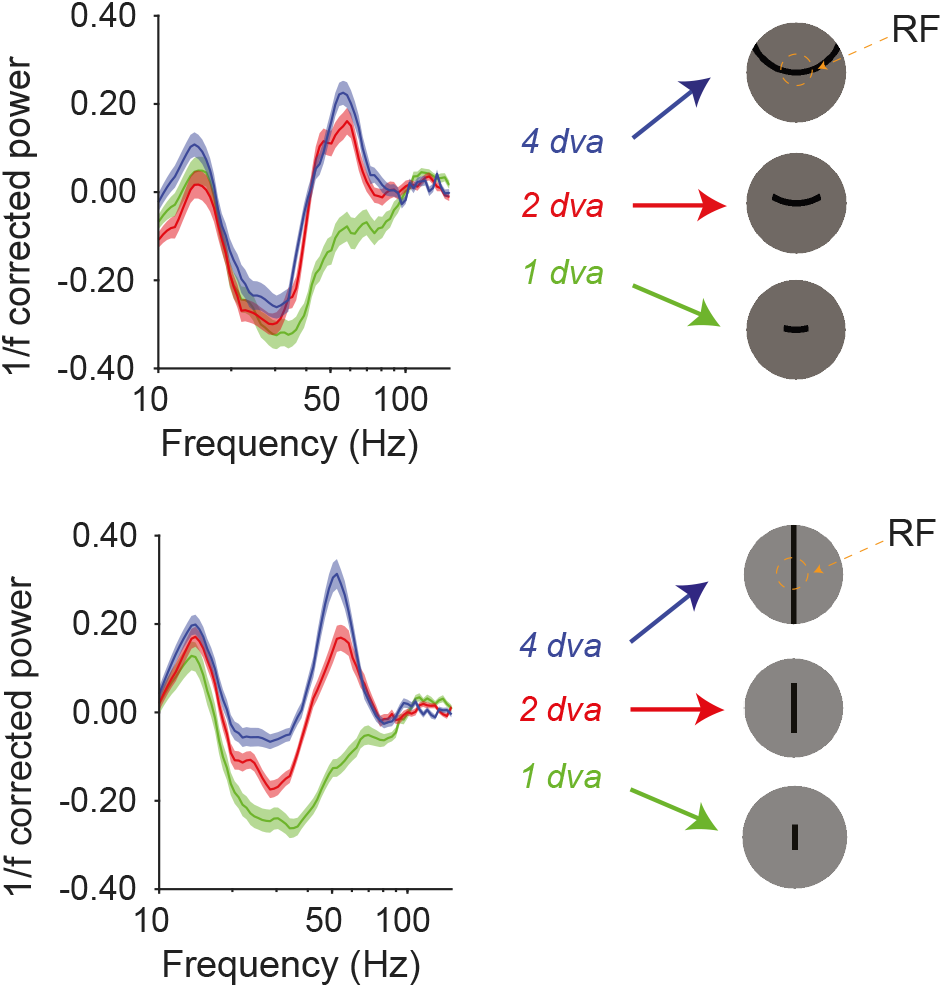
Curved bar (1 session, average across 14 channels) and straight bar stimuli (4 sessions, average across 53 channels) generate *γ* responses with characteristic size tuning beyond 1dva.

**Figure Extended Data 15:**
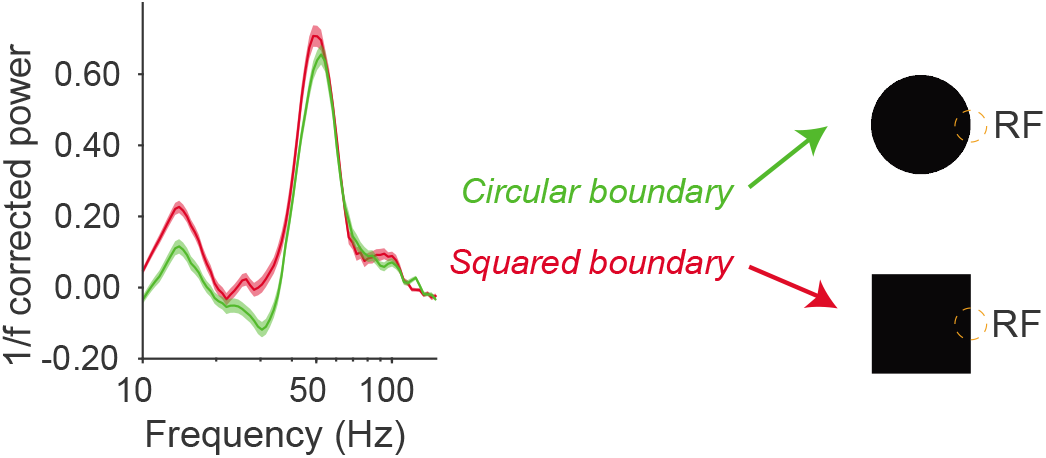
Boundary stimuli generate strong *γ* responses, both for curves and square filled surface stimuli (average across 14 channels, 1 session).

**Figure Extended Data 16:**
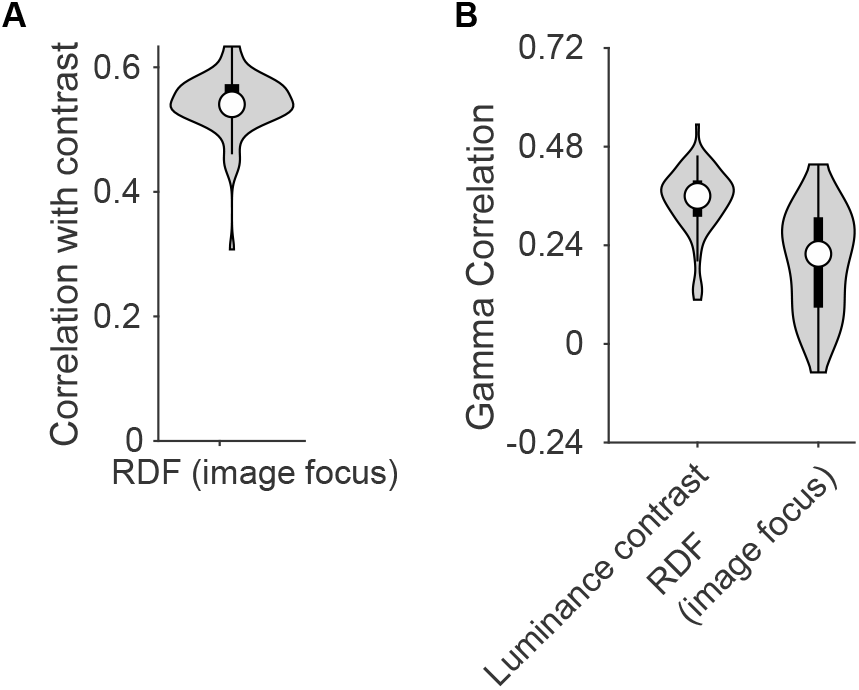
**(A)** Spearman correlation between image focus and luminance-contrast. **(B)** Correlation between *γ* and luminance-contrast or focus. Correlation of gamma with luminance-contrast was significantly higher than for image focus (paired T-test, *p* < 0.001).

**Figure Extended Data 17:**
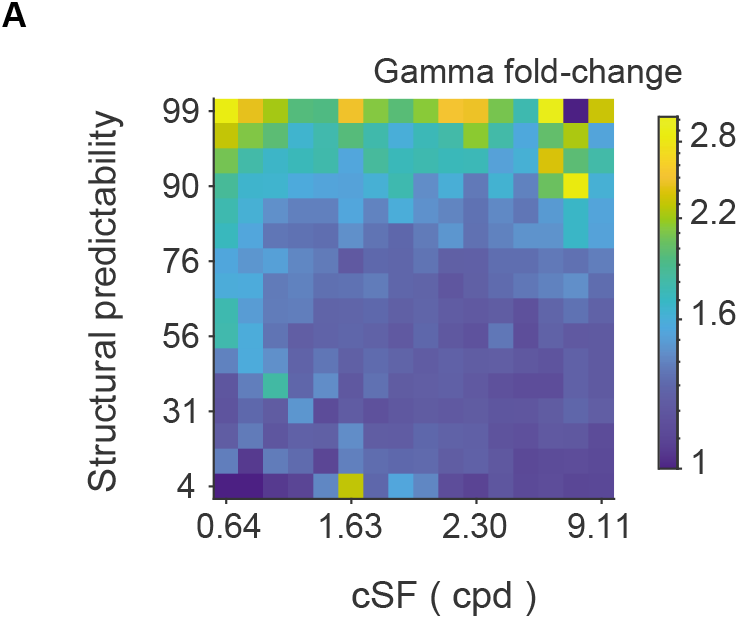
Gamma-band activity (fold-change) as a function of different values of spatial frequency in the RF and structural predictability. Predictability increases gamma across a wide range of spatial frequency values.

**Figure Extended Data 18:**
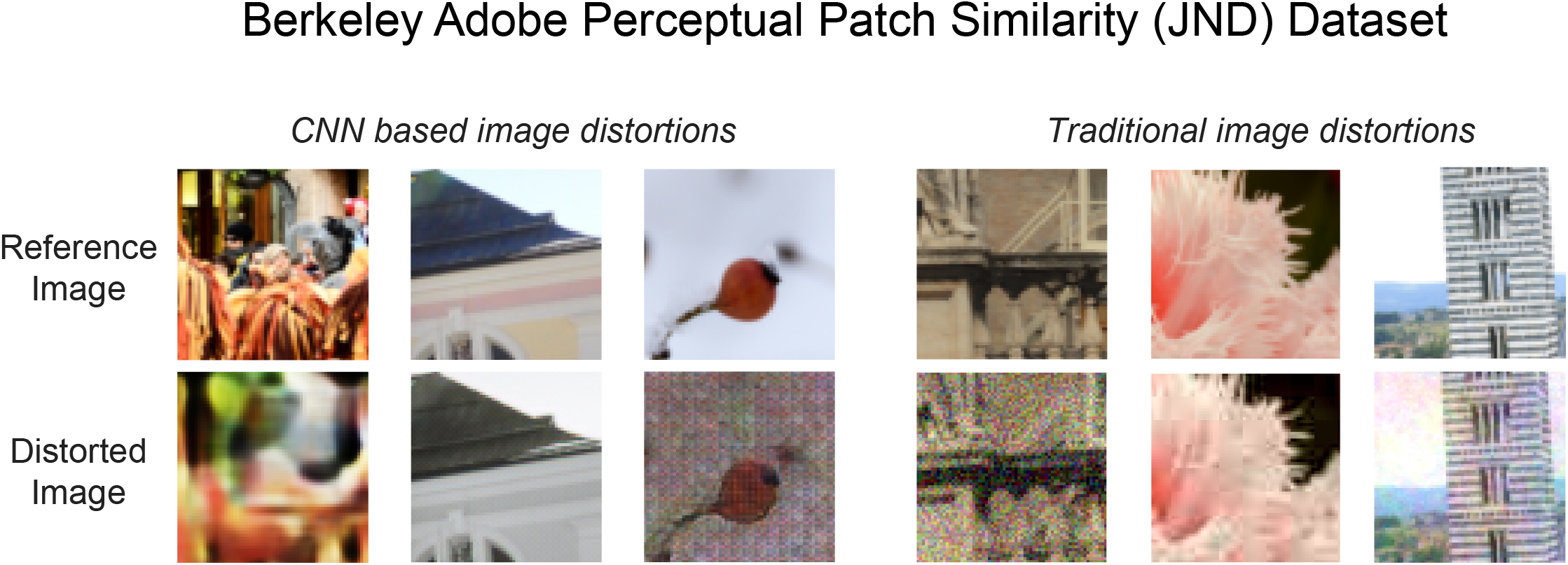
Examples of standard distorted images used to test perceptual similarity in humans.

**Figure Extended Data 19:**
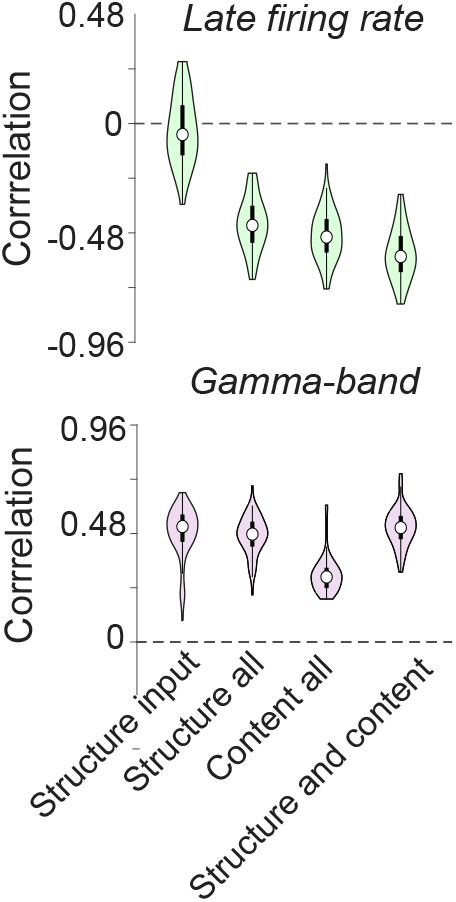
Left to right: (1) Correlation with structural predictability at the input image level; (2) regression model based on structural predictability across layers 1conv2, 2conv2, 3conv3, 4conv3, 5conv3; (3) regression based on content predictability in those layers; (4) regression based on both content and structure predictability in those 5 layers. For *γ*, the full regression model did not explain more variance than structural predictability at the input level (*P* = 0.40), and structure explained more variance than content (*P* < 0.001, paired T-test). For rates, the model with content explained more variance than structure (*P* < 0.001), and the model with both content and structure explained more variance than either content or structure (*P* < 0.001 for both).

**Figure Extended Data 20:**
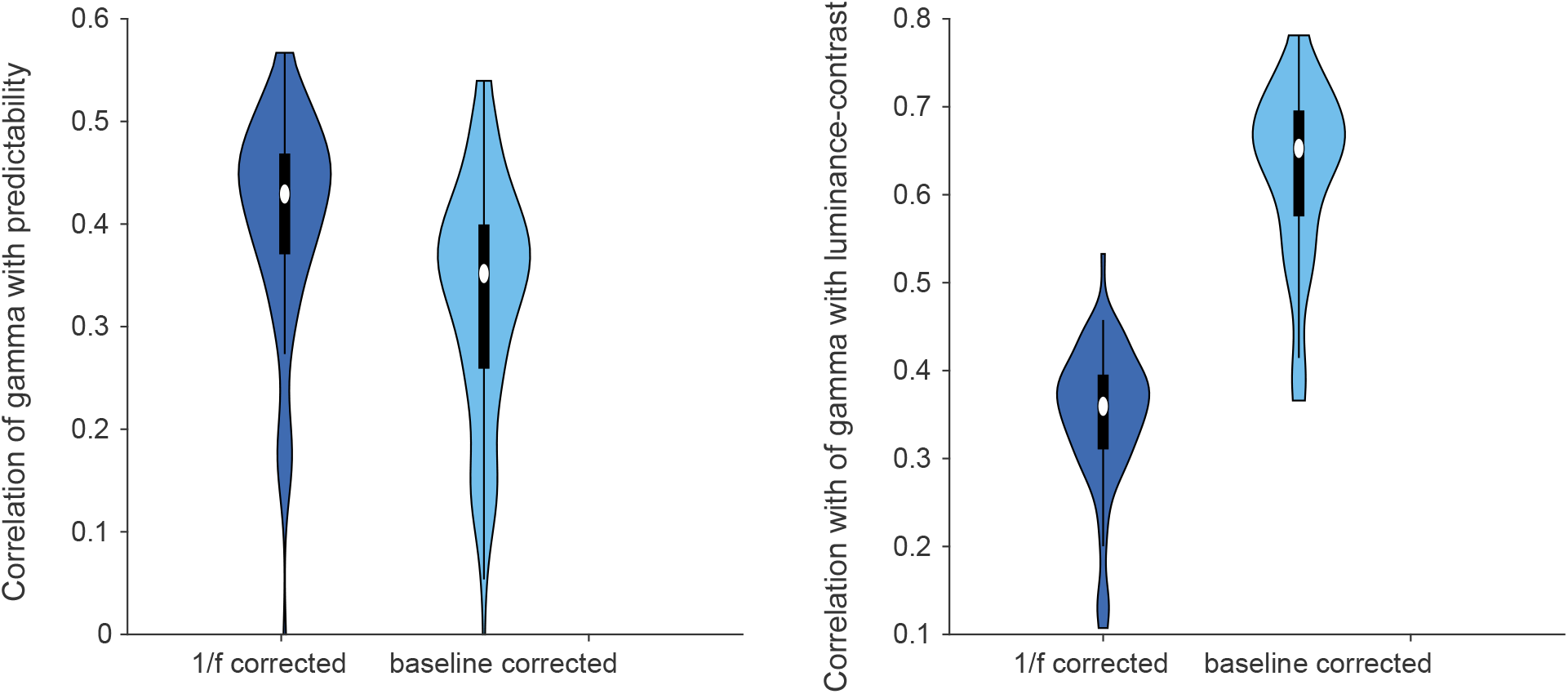
Correlation of *γ* derived either with 1/ *f* correction or as the baseline-corrected power. Predictability correlates more strongly with the 1/ *f* derived power, whereas contrast correlates much more strongly with the baseline-corrected power (*P* < 0.001 for both) which most likely reflects the influence of firing rates on baseline-corrected power.

**Figure Extended Data 21:**
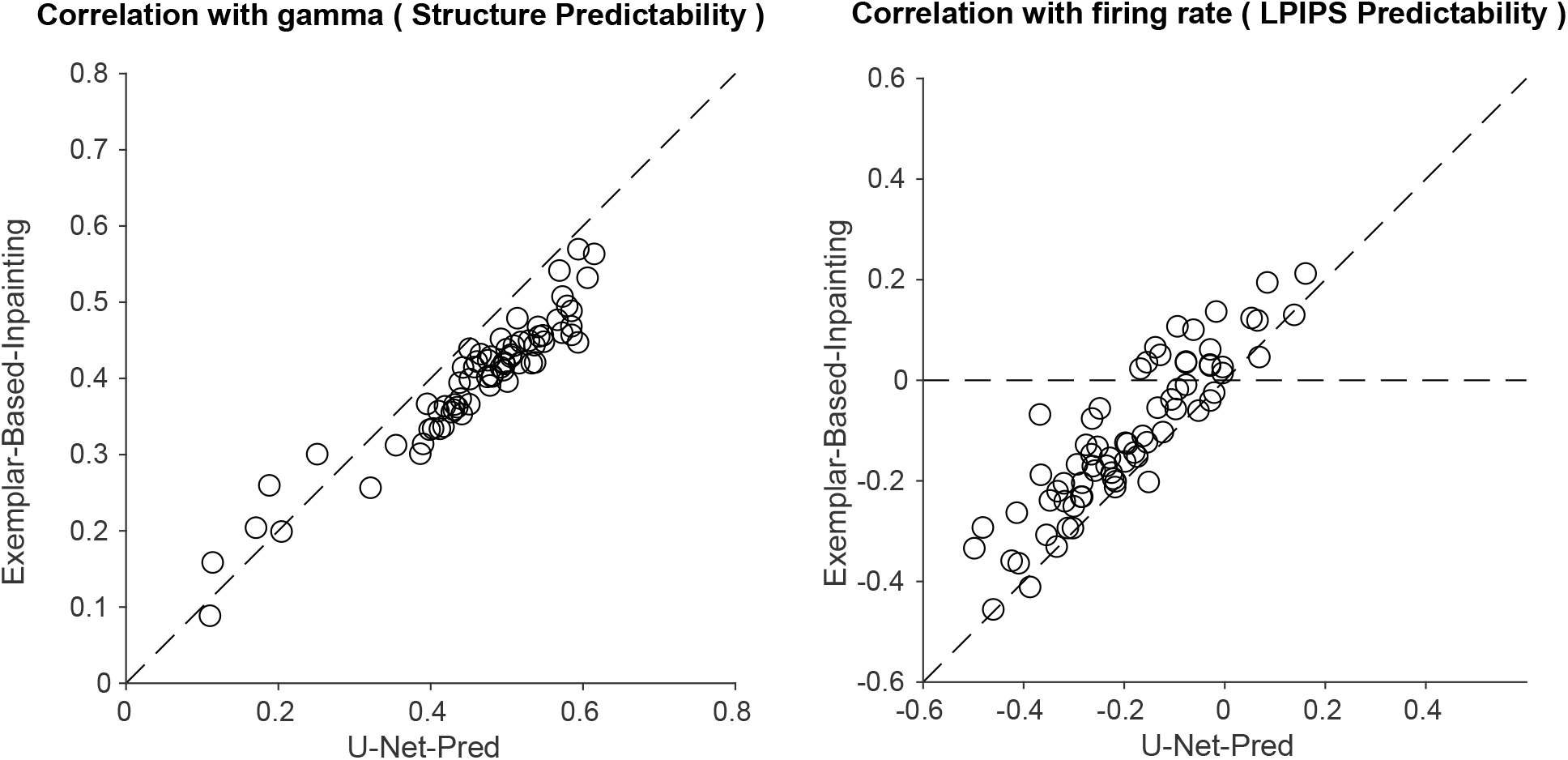
Comparison between predictability derived via deep learning methods and exemplar-based inpainting. Instead of making image predictions with a deep neural network with U-net architecture, we generated predictions using exemplar-based inpainting (107). Similar to the main analyses, we then computed predictability either as structural predictability or LPIPS-predictability. Predictability derived via exemplar-based inpainting correlated more poorly with *γ* for the structural predictability, but also shows a clear positive correlation with *γ* across channels. For firing rates, LPIPS correlations are stronger with deep learning methods (U-Net) than with exemplar-based inpainting.

